# Sequence constraints predispose Class D GPCRs to follow an atypical activation mechanism

**DOI:** 10.64898/2026.01.14.699465

**Authors:** Prateek Bansal, Diwakar Shukla

## Abstract

The biophysical principles underlying distinct conformational changes in proteins with similar topologies remain poorly understood. Class D G Protein-Coupled Receptors (GPCRs), fungal pheromone-sensing receptors essential for mating and survival, exhibit an atypical activation mechanism compared to other GPCR classes. Unlike Class A GPCRs, which activate through outward movement of TM6, Class D receptors undergo activation via outward displacement of TM7 coupled with inward movement of TM6. To investigate the origin of this atypical process, we employed state-specific generative AI sequence models to design protein sequences corresponding to unique active states, revealing that sequence constraints predispose Class D GPCRs toward this mechanism. We further explored the dynamic basis of activation through millisecond-scale atomistic simulations of STE2, a representative Class D GPCR and therapeutic target for fungal diseases. Using Maximum Entropy VAMPNets, an active learning based adaptive sampling strategy, we efficiently mapped the conformational free energy landscape of STE2. These simulations uncovered multiple intermediate states that have not yet been resolved experimentally and demonstrated that activation in these dimeric proteins involves fully decoupled monomers. Comparative simulations across Classes A, B, D, and F, totaling 4 milliseconds of all-atom molecular dynamics, revealed distinct patterns of electrostatic interactions. In Class D GPCRs, activation disrupts a dense TM7 interaction network while inward TM6 movement enhances electrostatic contacts. In contrast, Class A GPCRs such as CB1R display the opposite trend. Together, these findings show that sequence differences in TM6 and TM7 underlie the unique activation mechanism of STE2, offering new insights into GPCR conformational diversity.

## 1 Introduction

G Protein-Coupled Receptors (GPCRs) are cell surface receptors that transduce signals across the cell membrane in response to external stimuli. This response usually involves conformational transitions to an ‘active’ state, enabling association of the GPCR with G Proteins on the intracellular end and further facilitating downstream signaling. GPCRs are involved in maintaining homeostasis across vital biochemical signaling pathways, making them important drug targets. Recent studies estimate that between 26%^1^ and 36%^2^ of all FDA-approved drugs target GPCRs. Based on sequence alignments, GPCRs are classified into distinct families, with Class A GPCRs being the largest and most extensively studied.^3^ In contrast, Class D GPCRs, a small family expressed exclusively in fungi, are involved in mediating fungal development, virulence, and survival.^4^ Hence, these proteins are considered crucial targets for fungal diseases in humans. The absence of expression of these proteins in mammalian cells makes them an ideal orthogonal target for developing antifungal drugs.^5,6^ Unlike prototypical Class A or B GPCRs, Class D receptors function as homodimers. Despite this divergence, these proteins show the prototypical heptahelical transmembrane fold, a hallmark of GPCRs. Despite their therapeutic relevance and structural similarities, the activation dynamics of these proteins remain poorly understood. Hence, there is a need to study these proteins from a dynamic viewpoint. Only five Class D GPCR structures are available for the fungal STE2 receptor, showing how sparse the structural landscape of these receptors remains relative to other classes.

STE2 is a canonical fungal GPCR that mediates sensing of peptide pheromone *α*-factor,^7^ inducing conjugation and subsequent fusion with the opposite mating type in fungi.^8,9^ These GPCRs are expressed in all major pathogenic fungal species,^10^ hence studying STE2 activation dynamics can provide beneficial insights into targeting these proteins for rational design of broad antifungal therapies. A high degree of sequence conservation among separate fungal species such as *S. Kluyveri*, *C. Albicans*, *T. globosa*, *P. membranifaciens* and *D. bruxellensis* (Fig. S1) suggests that insights gained from this study can further be extended to other species. Activation dynamics of STE2 can further help elucidate the molecular basis of fungal signaling.

Recently, there has been an increased interest in the structural biology of STE2, with five structures resolved in the last five years. Structures of STE2 bound to *α*-factor have shed light onto the ligand recognition and overall structural changes occurring during ligand-induced activation.^11,12^ The G protein-bound active complex showed similarities to G protein-bound Class A GPCRs, with the notable exception of TM4 (TransMembrane helix 4).^12^ Further, this structure unveiled an entirely novel GPCR activation mechanism mediated by the outward movement of TM7, in contrast to canonical Class A/B1/F GPCR activation, which involves outward movement of TM6 instead.^12^ The dimeric interface involves an extensive interaction network between the protomers, and does not undergo any major conformational change during activation, in contrast to Class C GPCRs, where the protomers undergo major rotations with respect to each other, at the protomeric interface. ^13^ Considering the uniqueness of this active structure, there is a need to understand the activation of this family of GPCRs. Here, we address this question by using *S. cerevisiae* STE2 as a model system to investigate the dynamics of activation from a computational standpoint. In particular, four structures of STE2 - Inactive, Antagonist bound (PDB 7QA8^11^), Inactive-like intermediate (PDB 7QBC^11^), Active-like intermediate (PDB 7QBI^11^), and Active, G Protein Bound (PDB 7AD3^12^) are being used as starting points for simulations (Fig. 1). On comparing these structures, the major structural changes lie between the inactive and active structures (RMSD ∼ 3.9 Å, Fig. 1a). The active-like and the inactive-like structures are similar to the active and inactive structures, respectively. On comparing the inactive and inactive-like structures, we observed minimal differences in conformation (RMSD ∼ 0.8 Å, Fig. 1c), similar to the active-like and active structures (RMSD ∼ 0.6 Å, Fig. 1d). These structural comparisons raise the possibility of the existence of currently unidentified intermediate states that are structurally distinct from the inactive and active states themselves. Identification of intermediate states can enable a conformation-specific drug design, as shown in previous studies and shed light on the unique activation mechanisms of Class D GPCRs.^14,15^

**Figure 1:**
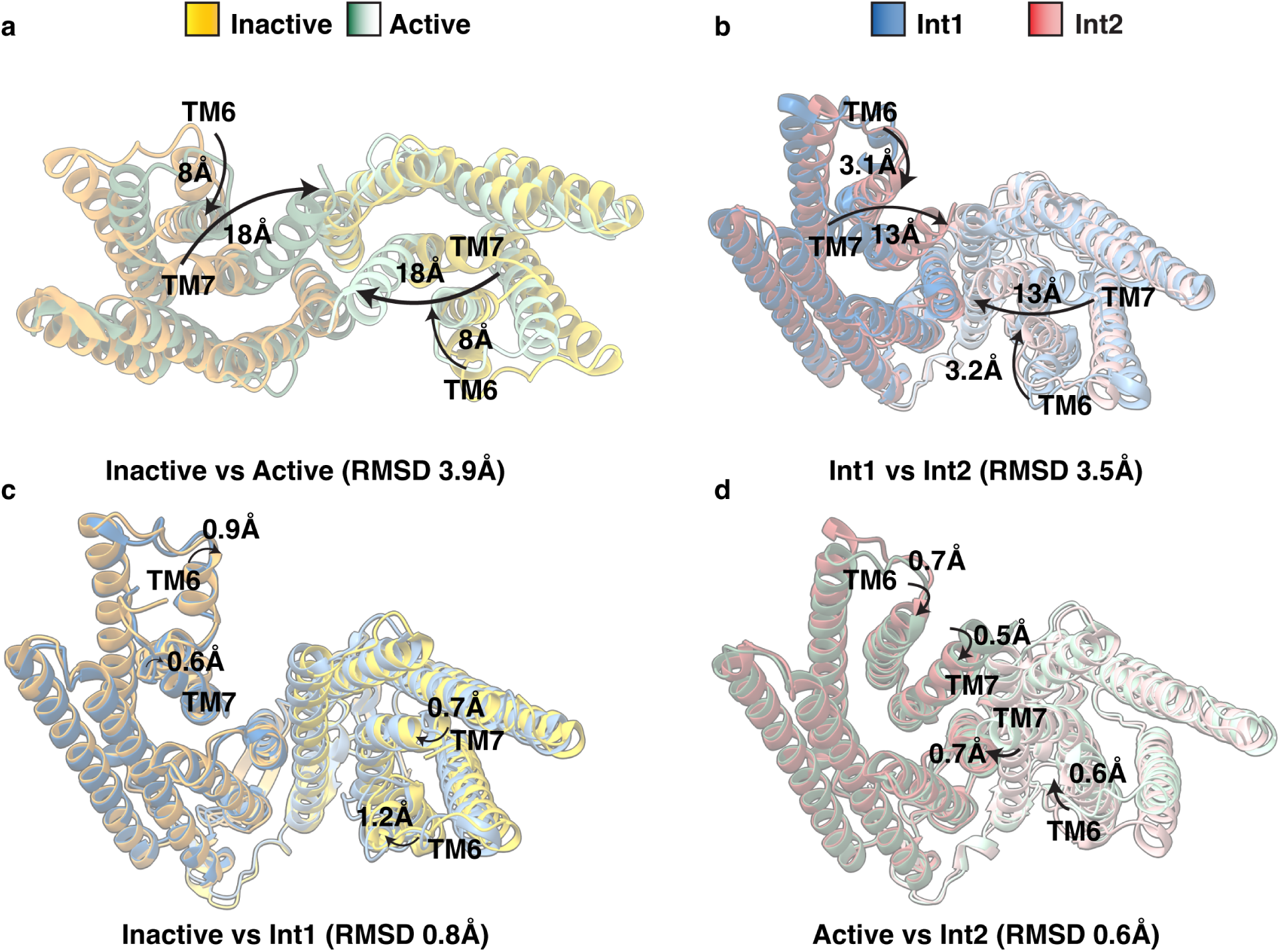
Comparison of structures of Class D GPCR STE2. Four resolved structures of are STE2 Inactive (PDB ID 7QA8^11^), STE2 Inactive-like (PDB ID 7QBC^11^), STE2 Active-like (PDB ID 7QBI^11^) and STE2 Active (PDB ID 7AD3^12^). (a) Structural changes between Inactive (7QA8^11^) and Active (7AD3^12^) STE2. Major structural changes are seen in TM7 and TM6. (b) Structural changes between Inactive-like (7QA8^11^) and Active-like (7AD3^12^) STE2. Major structural changes are seen in TM7 and TM6. Note that major differences observed in (a) are also reflected in (b). (c,d) Minor structural changes between Inactive (7QA8^11^) and Inactive-like (7QBC^11^) (c) and Active (7AD3^12^) and Active-like (7QBI^11^) (d). These structures show that the inactive-like and active-like intermediates are structurally similar to the inactive and active conformations, respectively, and raise the possibility of unresolved intermediate states that are structurally distinct from the active and inactive states.

We use a combination of machine learning, molecular dynamics simulations, and Markov State Models (MSMs)^16–20^ to obtain a complete picture of the activation mechanism from a thermodynamic and kinetic point of view. MSMs^14,21^ as well as long timescale MD^22,23^ have been recently used to successfully model GPCR function for non-class A GPCRs,^24–26^ including atypical activation.^27^ We first use ProteinMPNN-generated sequences conditioned on active backbones of Class D and A GPCRs to explain residue-level constraints that pre-dispose STE2 to follow an atypical activation mechanism. Using this as a motivation, we hence perform MD simulations using an adaptive sampling scheme ^28–30^ to sample the entire activation landscape from the inactive to active states, and show that activation occurs with a barrier of 4.2 ± 0.2 kcal/mol. The high dimensionality of the sampled data was reduced using time-lagged Independent Component Analysis (tICA),^31^ which projected the dataset onto dimensions that represented the slowest process being modeled using simulations. We show that the activation process is linked to the outward movement of TM7, in agreement with experiments. Using the constructed MSM and transition path theory, we resolve the millisecond-scale activation timescales of STE2 and identify novel intermediate states previously unobserved in experiments, and characterize residue-level movements that are unique to these states. We compare the activation process of STE2 with other classes of GPCRs, and show the residue level changes that enable Class D GPCRs to follow an atypical activation mechanism.

## 2 Results and Discussion

### 2.1 Sequence predisposes STE2 to follow an atypical activation mechanism

To investigate the cause of an atypical activation mechanism for Class D GPCRs, we sought to examine the hypothesis that the primary sequence controls the conformational constraints that allow the proteins to undergo activation. In particular, we asked which residues encoded in the STE2 sequence preferentially allow conformational changes centered on TM7 rather than the TM6-dominated motions typical of Class A GPCRs. Class A and Class D receptors both share the heptahelical transmembrane fold, so the sequential constraints imposed on either protein must differ for the proteins to undergo distinct activation mechanisms. To evaluate this hypothesis, we employed ProteinMPNN,^32^ a structure-conditioned, probabilistic sequence-design model. ProteinMPNN provides position-specific preferences consistent with a given backbone conformation, enabling an assessment of whether the native topology of STE2 allows for more sequential constraints on TM7 rather than TM6. We used these model-derived preferences to identify sites whose tolerated residue identities differ systematically from those in representative Class A sequences.

We specifically compared the sequences generated for the TM6 and TM7 helices for Class D receptor STE2 v/s a representative Class A receptor, CB1. Backbones of the active conformations were used as input topologies to the model, and the model was asked to generate 50000 sequences that could model the active conformation for both proteins. The most conserved residues along each helix were constrained. In addition, residues at the end of TM7 for the CB1 protein were held constant - corresponding to the NPxxY motif in Class A. We then calculated a consensus sequence for each helix by taking the residues that showed the highest frequency for each position. To compute the variability within the generated sequences and measure the model’s uncertainty associated with each position, we calculated the Shannon entropy associated with the probability distribution for each position. (Fig. 2). The shannon entropy plots for STE2 for sequences that model TM6 and TM7 (Fig. 2a, b) clearly show that the entropies associated with each residue position in TM6(Fig. 2a, orange highlight) are higher than the entropies associated with each position in TM7 (Fig. 2b, blue highlight). The high Shannon entropy associated with residues in TM6 can be interpreted as a measure of higher tolerance for sequence variability in STE2.

**Figure 2:**
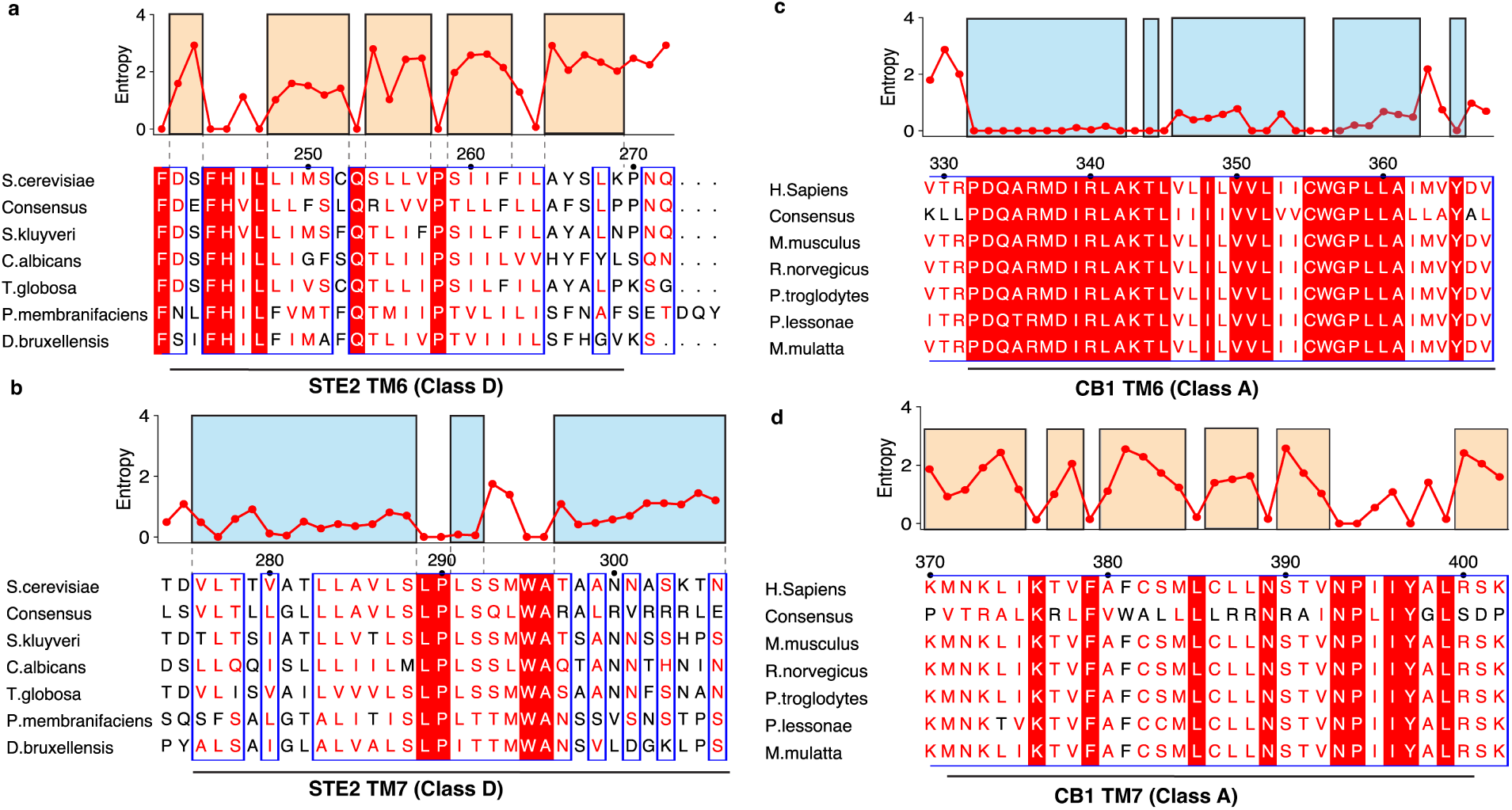
Sequence-encoded constraints differentiate activation helices in Class D and Class A GPCRs. (a, b) Shannon entropy per residue for ProteinMPNN-generated designs (50,000 sequences) conditioned on the active-state backbone of STE2 (Class D). TM6 (a, orange) exhibits higher entropy than TM7 (b, blue), indicating greater sequence tolerance at TM6 and tighter constraints on TM7. (c, d) Corresponding analysis for CB1 (Class A) conditioned on its active-state backbone shows the opposite trend: TM6 (c, blue) has lower entropy than TM7 (d, orange), consistent with tighter sequence constraints on TM6. For each helix, positions corresponding to the most conserved residues were fixed during design. Lower entropy reflects stronger sequence constraints and aligns with the helix primarily implicated in activation (TM7 in STE2, TM6 in CB1).

In contrast, the Shannon entropy associated with each position for TM6 and TM7 in CB1 showed the opposite trend - the sequences generated showed a lower Shannon entropy for residues associated with TM6 (Fig. 2c, blue highlight), as compared to TM7 (Fig. 2d, orange highlight). In addition, the model was also able to identify the most conserved residues for helix 6 as the consensus, even though the positions were not restrained during generation (Fig. 2c, blue highlight). With regard to the NPxxY motif, the model generated sequences with a low entropy for these positions, explaining the importance of these residues in the activation of Class A GPCRs. This further underscores the model’s reliability and its ability to act as a reliable proxy for computing the sequence tolerance associated with helices.

Collectively, these results support a model in which sequence-encoded constraints bias STE2 toward TM7-centered activation, in contrast to the TM6-dominated mechanism of CB1. Hence, we offer a sequence-level rationale for the atypical activation of Class D GPCRs.

### 2.2 Activation of STE2 involves blockade by TM7

Unlike canonical Class A/B1/F GPCRs, activation of class D GPCRs has been linked to the outward movement of TM7 and inward movement of TM6. However, the mechanistic details of this transition remain unclear. Hence, we performed millisecond-scale molecular dynamics simulations using an adaptive sampling strategy and constructed an MSM to recover the underlying thermodynamic and kinetic details of activation. To demonstrate this, we plotted the activation landscape of STE2 by taking all the simulation data, calculating the distance between V153^3.53^ (TM3) and A298^7.58^ (TM7) - to characterize the outward movement of TM7, as well as the distance between V153^3.53^ (TM3) and H245^6.37^ (TM6) - to characterize the inward movement of TM6 (Superscripts denote the numbering system introduced in Velazhahan et al. ^12^ specific to Class D GPCRs, analogous to the Ballesteros Weinstein numbering used for Class A GPCRs^33^). These distances were plotted against each other (Fig. 3a, Fig. S2a). In Fig. 3a, we can clearly observe that the major barrier associated with activation is the outward movement of TM7. These distances, when computed for the four PDB structures resolved for STE2, show that the inactive and inactive-like structures are located in the free-energy minima associated with an inward TM7 and outward TM6. On the contrary, the active-like and active structures lie in the minima associated with an outward TM7 and inward TM6. The barrier associated with activation is highest along the y-axis (4.1 ± 0.2 kcal/mol), showing that the outward movement of TM7 is the slowest process observed. Upon projecting the entire dataset along dimensions that model the slowest kinetic process observed in simulations (tIC1 and tIC2), we observe that the barrier associated with activation is 4.2 ± 0.2 kcal/mol (Fig. S3). To confirm that the slowest process is correlated with the outward movement of TM7, we projected the simulation data along the two tIC dimensions (Fig. 3b).

**Figure 3:**
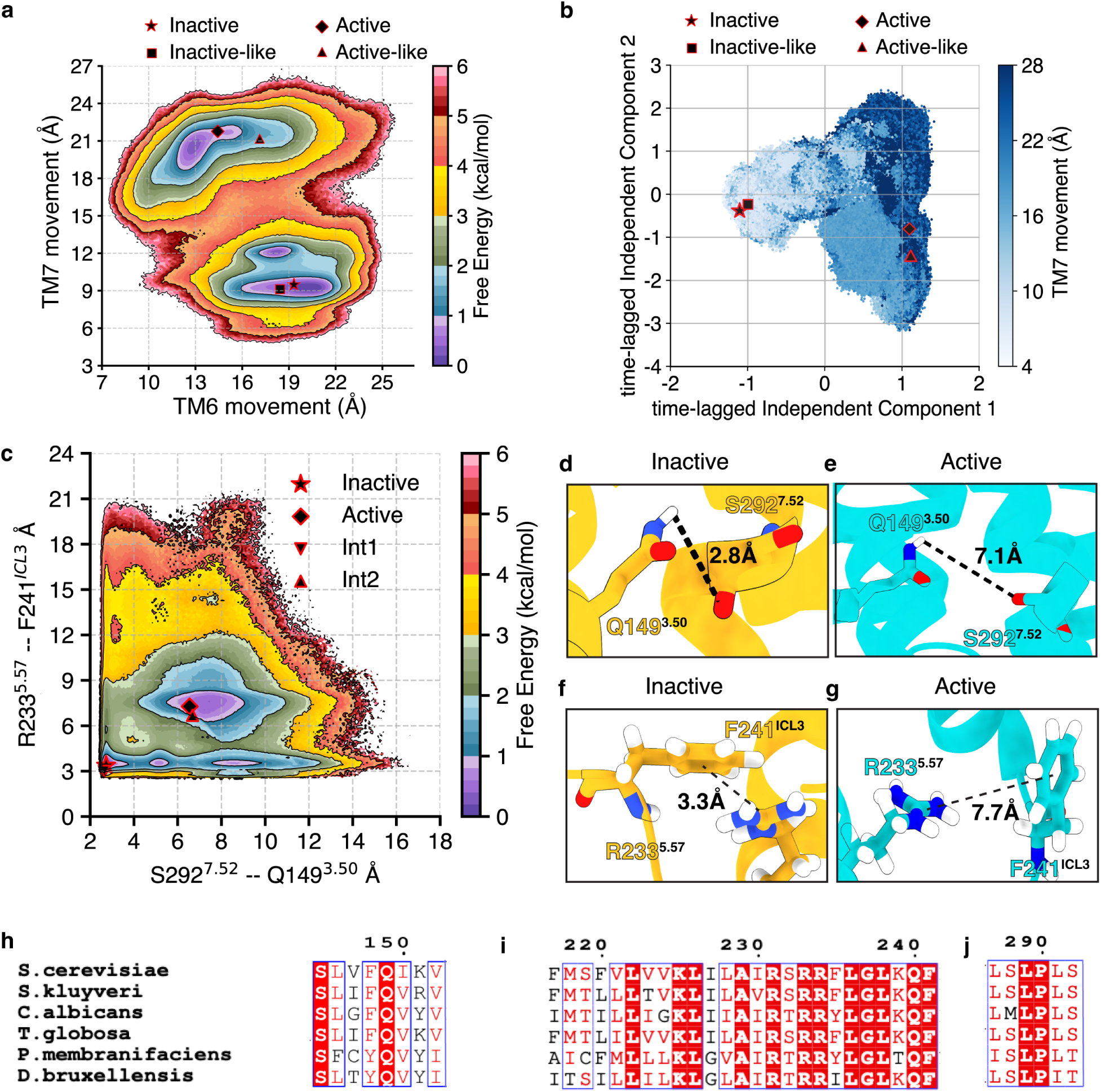
Outward movement of TM7 on STE2 activation. (a) The outward movement of TM7 plotted against the outward movement in TM6. (b) Scatter plot showing the outward movement of TM7 (colorbar) plotted along the two kinetically slowest processes (tIC1) and (tIC2). The gradient shows that the outward movement of TM7 is correlated with the slowest process observed in simulations. (c) The outward movement of TM7 plotted along with the cation-pi lock breakage seen in TM5. (d-g) Snapshots from simulation frames showing the breakage of the hydrogen bond between S292^7.52^ and Q149^3.50^ (top) and the cation-pi lock between F241*^ICL^*^3^ and R233^5.57^. Inactive state is shown in yellow, while active state is shown in cyan. (h-j) Multiple sequence alignments showing the conservation of Q149^3.50^, F241*^ICL^*^3^, R233^5.57^ and S292^7.52^. Complete sequence alignment in Fig. S1.

To further characterize residue level movements that occur on the outward movement of TM7, we computed the distance between S292^7.52^ and Q149^3.50^. This distance was plotted against the distance between F241*^ICL^*^3^ and R235^5.57^, a residue pair that shows the breakage of a cation-*π* lock on activation (Fig 2c). Upon calculating the free energy associated with each simulation frame and projecting the dataset along these two distances, we observe a barrier of ∼2.5 kcal/mol associated with the breaking of the cation-pi interaction, and a barrier of 1.8 kcal/mol associated with the breaking of the hydrogen bond (Fig. 3c, Fig. S2b). The subscripts indicate the identity of the protomer (A/B), similar to the jargon introduced in Velazhahan et al. ^12^ The free energy landscape shows a hydrogen bond in the inactive state between the amidic hydrogen of Q149^3.50^ and the alcoholic oxygen of S292^7.52^ (Fig. 3d). Upon activation, this hydrogen bond is broken, due to the outward movement of TM7 (Fig. 3e). This outward movement opens a cavity on the intracellular end of STE2 to accommodate the G protein (Fig. S4). This is further facilitated by the inward movement of TM6 (Fig. S4). This inward movement of TM6 leads to the breakage of the cation-pi lock in ICL3 (Fig. 3f,g). Here, we see that the pathway followed by the overall transition process is supported by the breakage of the hydrogen bond in TM7, followed by the breakage of the cation-pi lock, since that pathway is associated with lower free energy barriers as compared to the opposite. Furthermore, these residues are conserved across species (Fig. 3h-j Fig. S1), where S292 forms a part of the highly conserved LPLSSMWA motif in Class D GPCRs, analogous to the NPxxY motif in Class A GPCRs.^11^

Since STE2 forms a homodimer, we also observe breakage of the same locks in both the protomers (Fig. S3). Thus, the activation process is symmetric, and the association with the G protein can be mediated through either protomer. This symmetric activation suggests that either protomer is equally competent to recruit and activate the G protein, potentially enhancing signaling robustness or efficiency. This further differentiates Class D GPCRs from Class C GPCRs, where the G protein complex is asymmetric. The symmetric activation we observe in both protomers raises an important question: are the conformational changes in one protomer coupled to those in its partner, or can they proceed independently? To investigate this, we used our molecular dynamics simulation data to provide further insights.

### 2.3 Dynamics of the STE2 protomers is decoupled

Motivated by the symmetry of the activation locks, we analyzed the dataset for interprotomer coupling. To further investigate this, we focused on a pair of residues between TM4 and TM5, which were associated with the inward movement of TM5 during activation. We plotted the distance between S170^4.50^ and M218^5.42^ - a pair of residues that form a weak hydrogen bond between the thioether sulfur of M218^5.42^ and the alcoholic hydrogen of S170^4.50^, and constructed a 2D free energy landscape (Fig. 4a, Fig. S2c). The resulting free energy plot shows that the thermodynamic barriers associated with the breakage of the hydrogen bond in protomer A, followed by protomer B, and vice versa, are small and similar (∼ 1.1 ± 0.2 kcal/mol in protomer A vs ∼ 1.2 ± 0.2 kcal/mol in protomer B). This suggests that the protomeric transitions in STE2 are decoupled - meaning that the hydrogen bond can be broken for either protomer independently.

**Figure 4:**
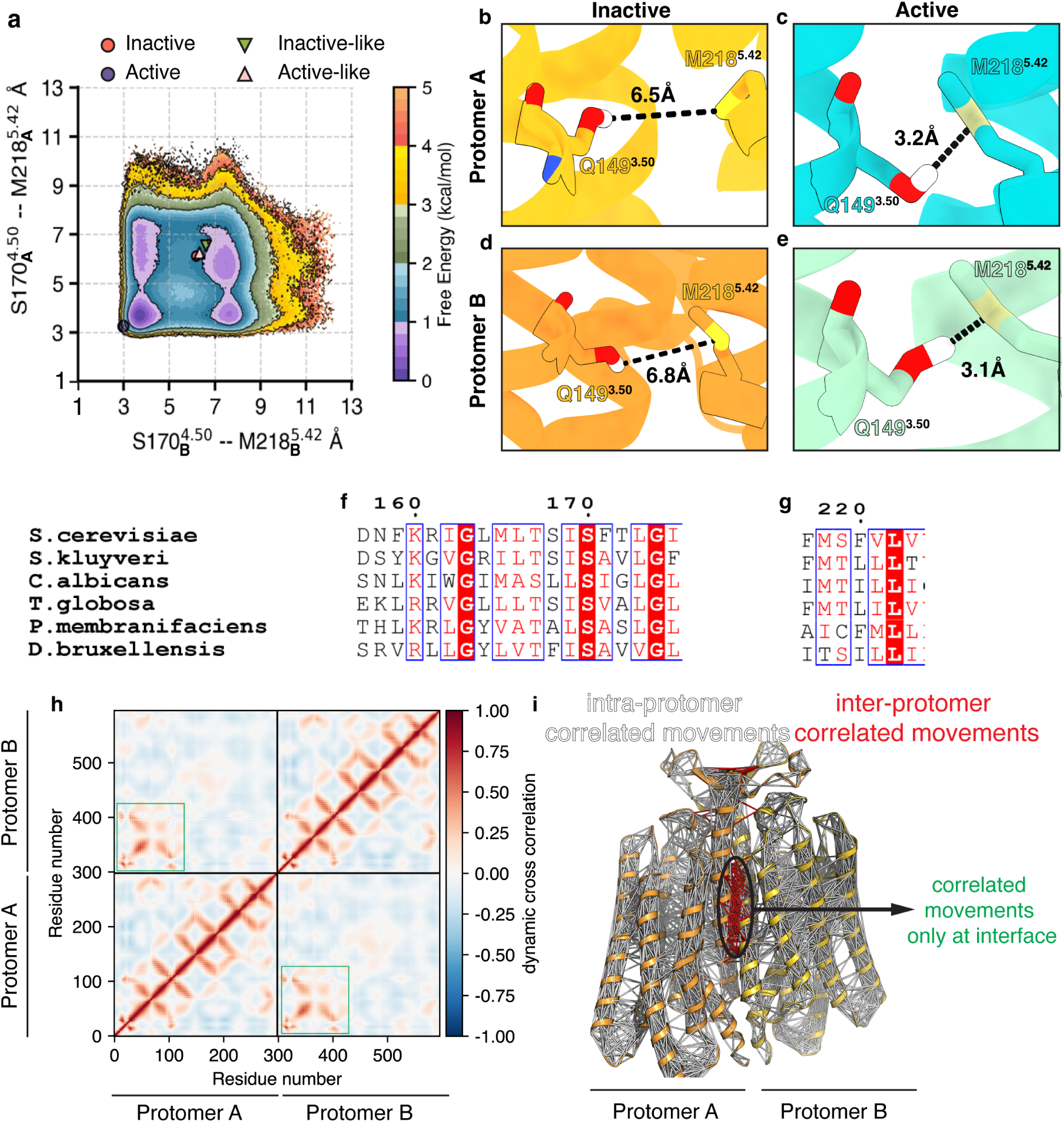
Dynamics of STE2 protomers is decoupled. (a) The inward movement of TM5 as a function of the free energy, plotted as a function of the distance between S170^4.50^ and M218^5.42^ for protomer A on the y-axis and protomer B on the x-axis. The inactive to active transition is associated with the formation of the hydrogen bond between these residues. (b-e) Snapshots from simulation frames showing the formation of the hydrogen bond between S170^4.50^ and M218^5.42^ for protomer A (b,c) and protomer B (d,e). The inactive state is associated with yellow (protomer A) and orange (protomer B), while the active state is associated with cyan (protomer A) and light sea green (protomer B) (f-g). Multiple sequence alignments showing the conservation of S170^4.50^ and M218^5.42^. (h) Dynamic cross-correlation for residue-level movements plotted as a heatmap. Inter-protomeric correlated movements are highlighted in green. (i) Dynamic cross-correlations shown on the backbone of STE2. The only coupling between the protomeric residue-level movements is observed at the interface. Intra-protomer correlated residue movements are represented as white sticks, while inter-protomer correlated residue movements are represented as red sticks.

To further support this decoupling, we used simulation snapshots - to identify intermediate states where the S170 - M218 hydrogen bond is broken in Protomer A but not Protomer B (Fig. 4b,c), and vice-versa (Fig. 4d,e). Interestingly, S170^4.50^ is the most conserved residue in TM4 across fungal species (Fig. 4f), while M218^5.42^ (Fig. 4g) also has a high conservation score. In the case of *D. bruxellensis*, M218^5.42^ is replaced by serine, another residue capable of forming side-chain hydrogen bonds with polar hydrogens. Since these residues are conserved, these insights can be generalized to other members of the family as well, providing crucial insights into the universal elements associated with the activation mechanisms of these proteins. We also computed the dynamic cross-correlations between residue movements for the entire dataset (Fig. 4h), showing that the residue movements within protomers show significant correlated movements, except at the protomeric interface (Fig. 4h, marked as a green box). This can be clearly observed when these correlations are projected on the protein backbone (Fig. 4i).

The decoupled activation of protomers also differentiates STE2 and other fungal GPCRs from canonical class C GPCRs, where the activation mechanism, being asymmetric, shows high coupling between the protomers. Yang et al. ^34^ have shown that for Class C GPCRs, the conformational changes in one protomer are highly coupled with the other protomer. From a biological context, we posit that this decoupling allows the cells to respond more sensitively to gradients in the concentration of the endogenous agonist *α*-factor. To further explain the decoupling of the protomers, we plotted the dataset presented in (Fig. 3a) but only when one of the protomers is active or inactive. For example, the dataset for protomer A was plotted when protomer B was completely active, and vice-versa (Fig. S5 a-c). On comparing the plots for the distances plotted for the entire dataset v/s the dataset with the plot when protomer B is active/inactive (Fig. S5 a-c), we observe a slight change in the free energy landscapes, but the minimas associated with the active state in protomer A are still present (Fig. S5b, c). A similar observation is made upon projecting the dataset on protomer B, but when protomer A is completely active/inactive (Fig. S5c-f). This shows that the activation of these proteins is completely decoupled and that the conformation of one protomer does not affect the conformational accessibility of the other protomer.

### 2.4 Transition between Intermediates 1 and 2 is the rate-determining step in activation of STE2

The intermediate states associated with STE2 activation that have been resolved experimentally do not show any major differences from their respective active and inactive states. Thus, in order to identify previously uncharacterized intermediates along the activation pathway, we used VAMPNet-based state space decomposition. VAMPnet is a deep learning method that learns non-linear projections of high-dimensional molecular dynamics trajectories optimized to preserve slow dynamical processes.^35^ Accordingly, the constructed model was able to identify three additional intermediate states that lie between the inactive and active states (named I1, I2, and I3), distinct from the inactive-like and active-like intermediate states reported in literature. The VAMPnet model identified the inactive-like and active-like intermediates as existing in the same macrostates as inactive and active macrostates, further corroborating that the resolved intermediates are kinetically and structurally close to the resolved inactive and active structures (Fig. S6). To characterize the transitions between the inactive and active states, we used the MSM and transition path theory^36^ to calculate reactive fluxes and timescales associated with the transition between the five states (Fig. 5a). These timescales can provide crucial insights into the activation process of STE2 and identify any potential bottlenecks in the overall activation process. To compute the average timescales associated with each transition, we defined the source (inactive) and destination (active) states by using the state decomposition from the VAMPnet model, and computed the mean first passage time, which is a measurement of the average timescales associated with the transition between the two states.

**Figure 5:**
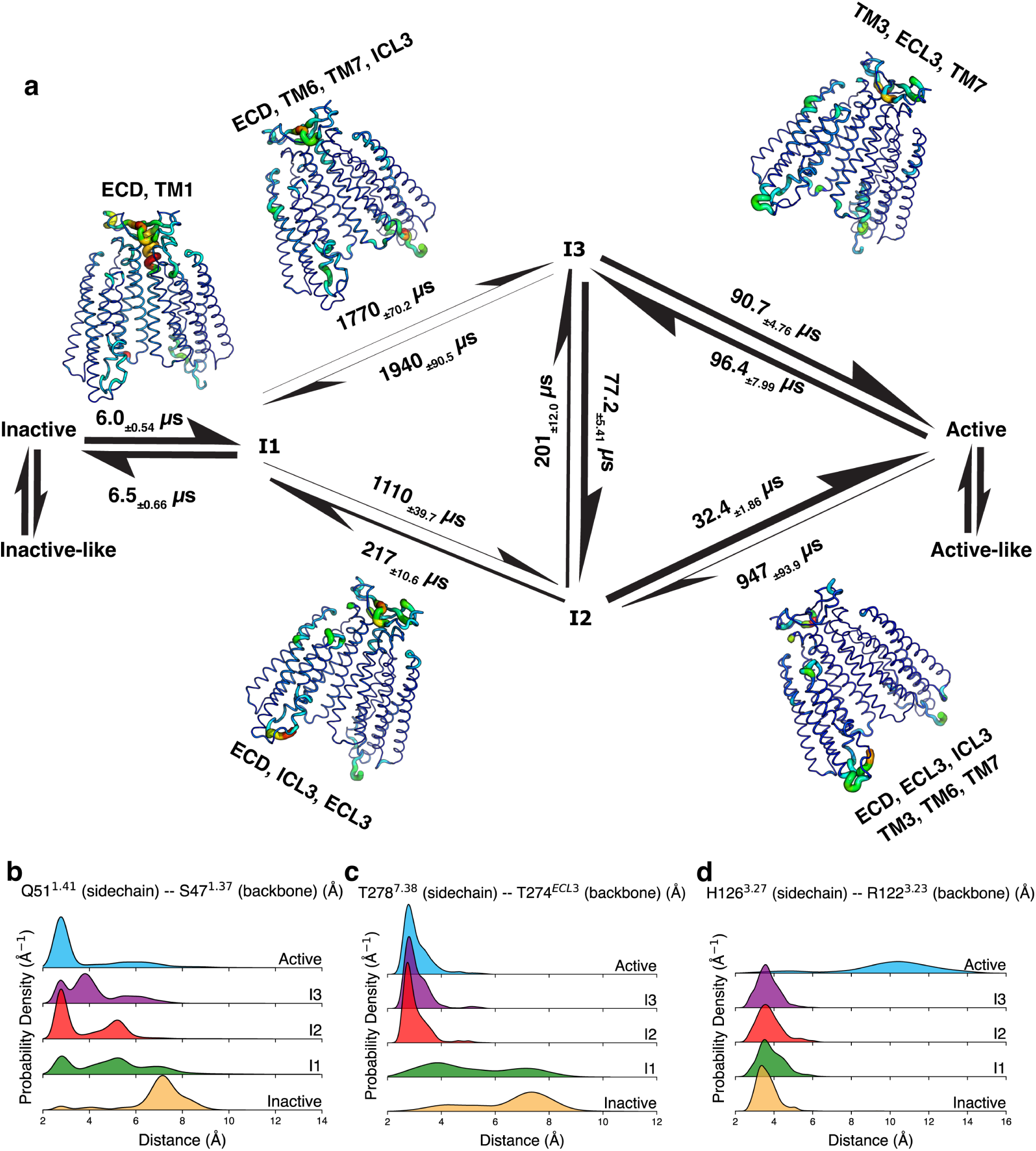
Activation pathways associated with STE2 activation. (a) Major structural transitions and timescales associated with each transition are separated with arrows. The thickness of arrows is proportional to the reactive flux between intermediate states. (b-d) Probability density plots for specific metrics that change on an intermediate-state basis. The probability density associated with each intermediate state is plotted separately. (b) plot for the distance between Q51^1.41^ and S47^1.37^ that forms a lock between inactive → I1 transition. (c) plot for the distance between T278^7.38^ and T274*^ECL^*^3^ that forms a lock between I1 → I2/I3 transition. (d) plot for the distance between H126^3.27^ and R122^3.23^ that breaks between I2/I3 → Active transition.

The first transition starting from the inactive state is the transition to intermediate state I1, occurring at a timescale of 6.0 ± 0.54 *µ*s. Major structural changes occurring during this transition include arrangements in the TM1 and ECD. From here, the transition from I1 to active is split into two pathways - one passing through I2 and the other through I3. The transition from I1 to I2 involves transitions in ECD, ICL3, and ECL3, while from I1 to I3 involves ECD, TM6, and ICL3. Here, the transition from I1 to I3 (1770 ± 70.2 *µ*s) is slower than I1 to I2 (1110 ± 39.7 *µ*s), owing to the inward movement of TM6 involved in I1 to I3 transition. The subsequent transitions from I2 and I3 to the active state involve the remaining structural changes, most importantly being the outward movement of TM7. The timescales associated with these transitions are in the sub-100*µ*s timescale (I3 to active, 90.7 ± 4.76 *µ*s, I2 to active, 32.4 ± 1.86 *µ*s).

To further establish the uniqueness of the identified novel intermediate states, we sought to identify molecular locks that were broken or formed during each transition along the activation pathway. To achieve this, 10000 frames from each macrostate were sampled according to the MSM probabilities, and an extensive interaction map was generated for each microstate, and interactions that broke or formed during each transition were identified (Fig. 5 b-d, Fig. S7, Fig. S8). During the first transition from Inactive → I1, conformational changes in TM1 are observed (Fig. 5a). This can be characterized by the formation of a hydrogen bond between Q51^1.41^ and S47^1.37^ (Fig. 5b, Fig. S7a,b). This is followed by the I1 → I2/I3 transition, which is characterized by the forming of a hydrogen bond between T278^7.38^ and T274*^ECL^*^3^ on the extracellular end of TM7 (Fig. 5c, Fig. S7c,d). Finally, the transition I2/I3 → Active is accompanied by the formation of a hydrogen bond between H126^3.27^ and R122^3.23^, stabilizing the active state (Fig. 5d, Fig. S7e,f). To distinguish I2 from I3, we found that the transition I2 → I3 involves the breakage of a hydrogen bond between N132^3.33^ and Q200*^ECL^*^2^ (Fig. S8a, b). Thus, these insights further underscore the uniqueness of the activation mechanism of Class D receptor STE2, and highlight the changes that occur as the protein transitions from inactive to active.

### 2.5 Comparison of activation mechanisms among classes shows uniqueness of Class D receptors

We have shown that the activation mechanism of STE2 is associated with the outward movement of TM7. This atypical mechanism sets it apart from other GPCR families, which primarily show activation associated with the outward movement of TM6. Previous studies have explored the activation mechanism of other GPCR families in atomistic detail.^37–40^ While previous experimental studies have compared activation mechanisms of different classes of GPCRs,^11^ they do so from a static point of view, only using cryo-EM structures. Since activation mechanisms are a strong function of the underlying conformational dynamics of the systems, here we compare the activation mechanisms of different classes of GPCRs from a dynamic viewpoint. We use the dataset generated in this study and compare it with previously published MD datasets.^38,39,41^ The associated free energy landscapes offer a dynamic perspective that delineates the activation process between inactive and active states. This enables the identification of metastable states, thermodynamic barriers, and alternate activation pathways not observable in cryo-EM structures alone. The free energy landscapes highlight mechanistic differences in activation across GPCR classes, and give us a way to identify the directionality of the movement of transmembrane helices, as well as establish which transmembrane helices are central to the activation process (Fig. 6a-h, Fig. S9a-c).

**Figure 6:**
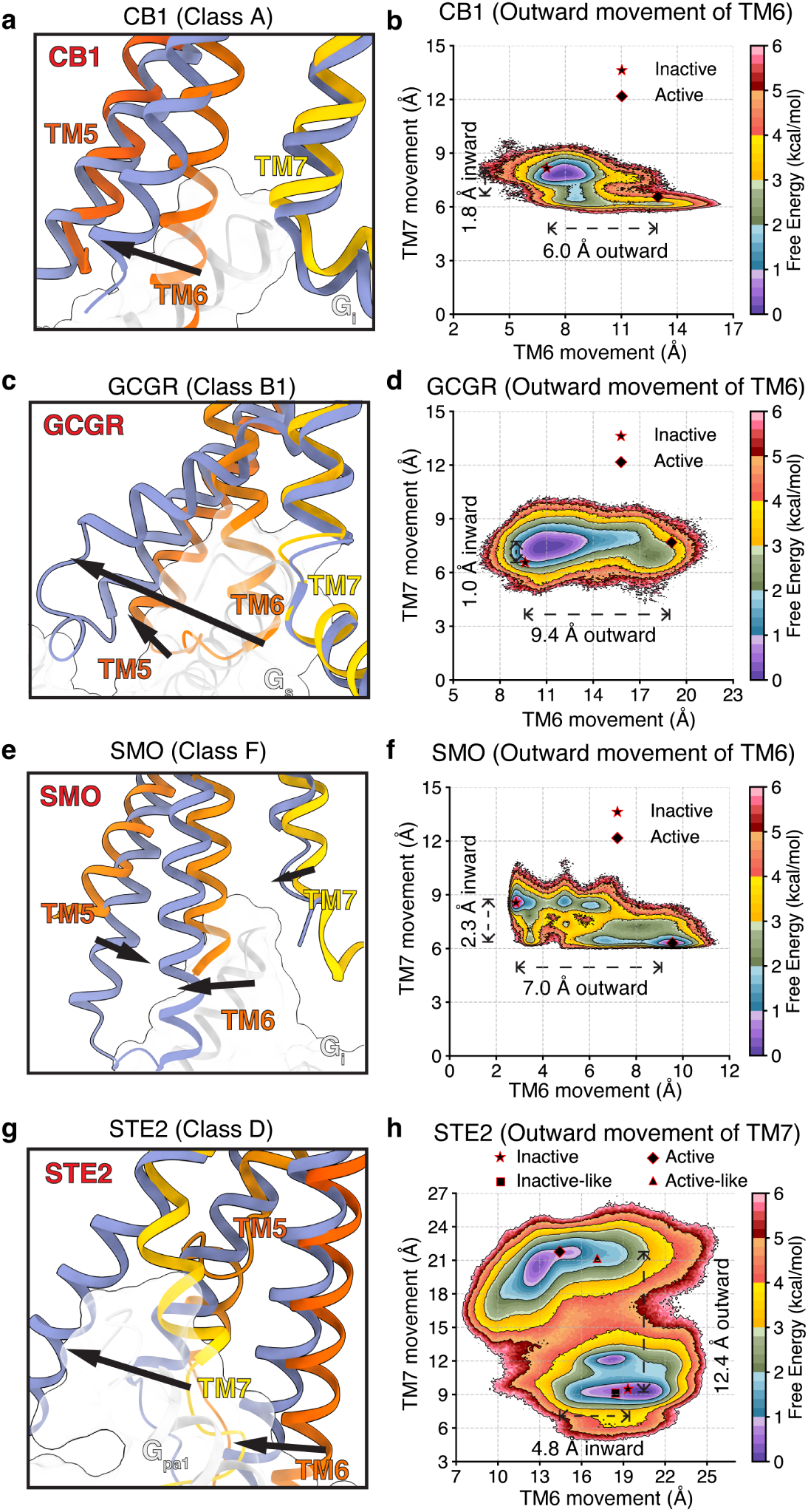
Comparison of activation mechanism across classes of GPCRs from a dynamic standpoint. (a) Snapshot from representative static resolved inactive and G protein bound structure for Class A receptor CB1 (PDB 5TGZ,^42^ 8GHV^43^), (b) Free energy landscape showing the activation mechanism involving the outward movement of TM6 in CB1 (c) Snapshot from representative static resolved inactive and G protein bound structure for Class B1 receptor GCGR (PDB 5YQZ,^44^ 6WPW^45^), (d) Free energy landscape showing the activation mechanism involving the outward movement of TM6 in GCGR (e) Snapshot from representative static resolved inactive and G protein bound structure for Class F receptor SMO (PDB 5L7D,^46^ 6XBL^47^), (f) Free energy landscape showing the activation mechanism involving the outward movement of TM6 in SMO (g) Snapshot from representative static resolved inactive and G protein bound structure for Class D receptor STE2 (PDB 7QA8,^11^ 7AD3^12^). (h) Free energy landscape showing the activation mechanism involving the outward movement of TM7 in STE2.

We use the following examples as representatives for each class of proteins - CB1 (Class A), GCGR (Class B1), SMO (Class F), and STE2 (Class D). For Class A receptor CB1 (Fig. 6a,b, Fig. S9a), a previous study has observed classical TM6 outward and TM7 inward movement as the chief activation mechanism.^41^ For Class B1 receptor GCGR (Fig. 6c,d, Fig. S9b), previous studies have studied TM6 kinking and outward motion as a unique feature associated with activation.^39,48^ For Class F receptor SMO, our previous study,^38^ shows a less dramatic TM6 shift, while TM5 plays a more prominent role in activation(Fig. 6e,f, Fig. S9c). However, the common element among all of these classes has been the outward movement of TM6. Each of these classes shows a 3-5 kcal/mol as the highest barrier associated with activation, and a sub-millisecond activation process.^38,39,41^ Here, in Class D receptor STE2, TM7 is the primary helix associated with outward movement, novel among GPCR classes(Fig. 6g,h, Fig. S2a).

Interestingly, the free energy barriers and the activation timescales remain in the same order of magnitude as other classes, while showcasing an entirely unobserved activation mechanism. These distinctions in activation mechanisms reflect an evolutionary divergence in signaling among GPCR classes, particularly as Class D receptors are unique to fungi and function as symmetric dimers. This gives us a unique opportunity to design orthogonal ligands that target Class D GPCRs in the case of fungal diseases, but further studies are needed to explore this proposition in greater detail.

On comparing the activation mechanisms for Class A GPCR CB1 and Class F GPCR SMO, the transition from inactive to active states passes through three intermediate states.^38,41^ On the other hand, Class B1 GPCRs are associated with a single major intermediate state. While Class A GPCRs are associated with ∼500*µ*s timescales, Class B1 GPCRs.^39^ On comparison with other classes of GPCRs, we observe that the transition timescales are similar to Class A and B1, with CB1 activation timescales in the milliseconds and GCGR activation timescales in hundreds of microseconds.^39^ Thus, our work identifies STE2 to have similar timescales of activation as other classes of GPCRs.

Finally, to further understand this observation from a structural point of view, we computed the interaction energies for inactive and active conformations of STE2 and CB1. Specifically, the electrostatic and Van der Waals interaction energies were calculated for the interactions of TM6 and TM7 with the rest of the protein (Fig. S10). The energies demonstrate that the outward movements of TM7 in STE2 and TM6 in CB1 are associated specifically with the decrease in electrostatic interactions (Fig. S10a, b), over Van der Waals interactions. In STE2, TM6’s interaction with the rest of the protein increases, leading to an increase in the electrostatic interaction energy (Fig. S10a). To further illustrate the role of TM7 for STE2, we computed the residue-wise contributions to the electrostatic energy for active and inactive conformations, and plotted the difference between interaction energies (Fig. S11a). The plot demonstrates that residues toward the intracellular end of TM7 contribute to the highest difference in interaction energies. This can be demonstrated by plotting the electrostatic networks in the inactive and active conformations (Fig. S11b,c). Residues that showed the greatest change in electrostatic energy were chosen (S288^7.48^, S292^7.52^, S293^7.53^, W295^7.55^, and N300^7.60^). The inactive conformation showed a dense electrostatic network involving TM3, TM5, and TM6 (Fig. S11b). Compared to the active conformation, the inactive state shows a sparse electrostatic network (Fig. S11c). The loss of electrostatic contacts in the inactive state (shown by positive values for S292^7.52^, S293^7.53^, W295^7.55^ and N300^7.60^ in Fig. S11a) is countered by the formation of a sparse network by S288^7.48^ (negative, Fig. S11a). These residues are also associated with sequence entropies (Fig. 2a), further validating the sequence-based constraints present in TM7 for STE2.

## 3 Conclusions

In this study, we provide a mechanistic understanding of the activation of a representative Class D fungal GPCR, STE2, using machine learning, molecular dynamics simulations, and Markov state models. We show that sequence-encoded constraints predispose STE2 to follow the atypical activation mechanism. We provide a detailed overview of the mechanistic process that occurs during activation of Class D GPCRs. We show that STE2 activation is associated with the outward displacement of TM7 and the inward movement of TM6 - a mechanism distinct from canonical GPCR classes, which involve the outward movement of TM6 to accommodate the G protein. The activation of STE2 occurs symmetrically in both protomers, and the protomers are decoupled from each other. Either protomer can show an active state, irrespective of the other protomer being active or inactive. This distinguishes the dimeric STE2 activation from the asymmetric activation observed in dimeric Class C GPCRs. To further dive into the details of the activation pathway, we utilized VAMPnets to discretize the entire ensemble into 5 macrostates - Inactive, I1, I2, I3, and Active. We studied the kinetics of these transitions and determined that the rate-determining step in the entire activation process is associated with the transition from intermediates I1 → I3, occurring on a millisecond timescale. The highest free energy barrier associated with the overall activation process is approximately 4.2 ± 0.2 kcal/mol. We also identified multiple molecular locks unique to the transition between each intermediate state, thus providing a residue-level overview of the activation process.

By comparing free energy landscapes across Class A (CB1), B1 (GCGR), F (SMO), and D (STE2) GPCRs, we highlight the differences as well as similarities associated with the activation mechanisms of these classes of proteins. To investigate the proclivity of Class D receptors like STE2 to undergo an atypical activation mechanism, we employed ProteinMPNN and generated sequences to show that sequence constraints predispose STE2 to follow an atypical activation mechanism. Through this study, we are able to provide an enhanced understanding of Class D receptor function. A high degree of sequence conservation among separate fungal species suggests that insights gained from this study can be further extended to other species. We posit that insights from this work will be useful for designing more selective antifungal therapies that target these proteins.

## 4 Methods

### 4.1 Molecular Dynamics Simulations

#### 4.1.1 System Preparation

For simulating the activation process of Class D receptor STE2, molecular dynamics simulations were performed using 4 different starting points - from the ligand-free apo state (PDB ID 7QB9^11^), Inactive-like Intermediate state (PDB ID 7QBC^11^), active-like intermediate state (PDB ID 7QBI^11^), and active state (PDB ID 7AD3^12^). For inactive-like intermediate state, active-like intermediate state system, and active state systems, the bound peptide agonists were removed. For the active state system, the bound G protein complex was removed. Missing residues were added to each protomer to ensure each protomer had the same number of residues in each system (S9 - N306). Protonations were checked using propka3, ^49^ and accordingly, none of the residues were protonated. Each of the 4 STE2 conformations was embedded in a lipid bilayer, and the composition of the bilayer was based on to plasma membrane composition of *Saccharomyces cerevisiae* ^50^(Table S1). Each protomer was capped with neutral termini ACE for the N-terminus and NME for the C-terminus, respectively. Each system was solvated using water molecules according to the TIP3P water model^51,52^ and 150 mM KCl,^53^ to mimic physiological conditions. Atomic interactions were modeled with the CHARMM36m force field.^54–57^ Each protein was embedded into the bilayer and solvated using CHARMM-GUI.^58–65^ System sizes were 87932 (Active STE2), 89960 (Inactive STE2), 90765 (Active-like intermediate), and 91693 (Inactive-like intermediate). Non-protein hydrogen masses were repartitioned to 3.024Da to enable usage of a longer timestep, 4fs. ^66^ The only difference between the 4 starting states was the conformation of the protein in the system.

#### 4.1.2 Pre-Production Simulations

Each system underwent energy minimization for 50000 steps. All hydrogens were constrained throughout all simulations using the SHAKE algorithm. The protein backbone was restrained during the pre-production steps, using a harmonic potential with spring constant 10 kcal/(mol.A^°2^). Each system was then heated from 0 to 310K over a course of 10 ns of simulations, with a linear increase in temperature, using the NVT ensemble. The Langevin thermostat was used for this purpose. The systems were then pressurized using the NPT ensemble and the MonteCarloMembraneBarostat for 10 ns. Then the system was equilibrated for 40 ns with restraints, and 40 ns without restraints, out of which the first 20 ns were discarded. All pre-production simulations were performed using OpenMM7.7^67^ on a NVIDIA RTX 4090 GPU on a local machine.

#### 4.1.3 Production Simulations

Production simulations were run using OpenMM 7.7 on GPUs donated by citizen scientists on the distributed computing project Folding@Home.^68^ Simulations were run in an iterative round-wise fashion for each system, where simulations for the first round were seeded from the equilibration trajectory, running hundreds of short trajectories in parallel for each system. Simulations after every round were collected and pooled, and new starting points for the next round of simulations were chosen using an adaptive sampling strategy to encourage exploration of the conformational space. Simulations were performed until the pooled data from all 4 systems were connected in the tIC space, with the first two components showing continuous density from the inactive to active starting points and vice versa.

### 4.2 Adaptive Sampling

To encourage conformational exploration and sampling rare transition states rather than stable states, an adaptive sampling strategy based on maximum entropy VAMPnets^28^ was used. Recently, many machine learning based adaptive sampling strategies have been developed that allow for efficient sampling of conformational dynamics of proteins.^29,69–72^ MaxEnt VAMPnet iteratively analyzes ongoing MD trajectories, identifies states with high uncertainty or entropy, and selectively launches new simulations to sample those regions. By maximizing Shannon entropy across predicted state distributions, the method ensures broad coverage of conformational space rather than overfitting to already well-sampled states. Simulations from each of the 4 systems at the end of each round were pooled, and collective variables (Table S2) were calculated for featurizing each frame. Frames that maximized the entropy of the state decomposition for the next round of simulations were chosen as the next starting points. The sampling strategy used the instantaneous and time-lagged datasets, both extracted from the simulation data collected until the previous round of simulations. The inputs were fed to a VAMPnet^35^ that was used for reducing dimensionality to a 6-dimensional latent space, yielding two latent spaces, corresponding to the instantaneous and time-lagged datasets. The VAMP2 score between the instantaneous and time-lagged latent spaces was used as the loss function. Once the neural net was trained, frames from the dataset that maximized the Shannon entropy, which is a measure of the error in the dimensional reduction of the system, were used. For further methodological details, please refer to Kleiman and Shukla, 2023 (ref.^28^). The data collected in each round is presented in Fig. S12-S14 and Table S3.

### 4.3 Dimensionality Reduction using tICA

To reduce the high-dimensional nature of the dataset, the trajectories were featurized, where each frame of simulation was represented as a row of 85 features. Time-lagged Independent Component Analysis (tICA) was used to obtain a linear combination of features, where the first dimension corresponded to the slowest process associated with the system (Fig. S3). The coefficients used for the linear combinations of the features were obtained by first computing the covariance between instantaneous and time-lagged datasets to maximize the time-lagged autocorrelations. This ensures that the dimensional representations correspond to the slowest dynamical processes observed in simulations.

### 4.4 Markov State Model Construction

Markov state models^16–20^ were constructed by performing a state decomposition using minibatch k-means clustering on the reduced-dimensional representations obtained from tICA. The microstates defined by the state decomposition were then subjected to counting, where inter-state counts were observed, which were at least a lag time (*τ*) apart. To optimize the lag time and maintain Markovianity across the model, implied timescales were calculated as a function of the lag time (Fig. S15a), and a lag time was chosen (30 ns) such that the implied timescales beyond the lag time were invariant with the lag time. To optimize the number of tIC dimensions used as input to the clustering algorithm, as well as the number of clusters (microstates) for the state decomposition, a grid-search approach was used (Fig. S15b). The optimization was performed by comparing the VAMP2 score, which measures the degree to which the chosen states and input dimensions are able to capture the slowest kinetic processes observed in simulations. Accordingly, 1200 clusters and 8 tIC dimensions were used for constructing the Markov model. Once constructed, the Markov model was validated using a 5-state Chapman-Kolmogorov test (Fig. S16). Once validated, the constructed MSM was used to reweigh the probabilities of each frame in the simulation, eliminating the sampling bias introduced using adaptive sampling. The MSM object was also used to compute the reactive fluxes necessary for computing the kinetic timescales associated with the conformational transitions.

### 4.5 ProteinMPNN for sequence generation

To generate sequences associated with protein backbones for Class D GPCR STE2 and Class A GPCR CB1, ProteinMPNN^32^ was used. For STE2, the active state backbone (PDB ID: 7AD3) was used as an input. Residues from each protomer were tied within the protein using the homooligomer option of the make_tied_positions_dict helper script, a part of the package. The residue-level constraints used to model the protein chains are reported in (Table S4). For CB1, the active state backbone (PDB ID: 8GHV) was used as input. The residue-level constraints used to model the protein chains are reported in (Table S5). For each protein, 50,000 sequences were generated. A sampling temperature of 0.2 was used. A batch size of 1 was used for generating the sequences. Once sequences were generated, the multi-sequence alignments were performed using in-house scripts, Biopython^73^ and Clustal-Omega.^74^

## 5 Acknowledgements

The authors would like to thank the volunteers from the Folding@Home distributed computing project for providing compute time for simulations. D.S. acknowledges support from NIH grant R35GM142745. P.B. acknowledges support from the A.T. Widiger Fellowship awarded by the Department of Chemical and Biomolecular Engineering at the University of Illinois.

## 6 Data and Code Availability

Codes used for analysis are available at https://github.com/ShuklaGroup/ClassD_Activation (GitHub). Trajectories and parameter files are available on Box: https://uofi.box.com/s/i4eoid4n6kumifgqw2a4pks55ghxek4y

