## Supplementary Information for "Sequence constraints predispose Class D GPCRs to follow an atypical activation mechanism"

### Supporting Information for: Sequence constraints predispose Class D GPCRs to follow an atypical activation mechanism

#### List of Figures

S1 Multiple Sequence Alignments . . . . . . . . . . . . . . . . . . . . . . . . . . S3
S2 Errors for plots in Fig 2 and Fig 3 . . . . . . . . . . . . . . . . . . . . . . . . S4
S3 TICAplot . . . . . . . . . . . . . . . . . . . . . . . . . . . . . . . . . . . . . S5
S4 Inward movement of TM6 . . . . . . . . . . . . . . . . . . . . . . . . . . . . S6
S5 Activation metrics for monomer-specific activation . . . . . . . . . . . . . . . . S7
S6 Macrostate Decomposition . . . . . . . . . . . . . . . . . . . . . . . . . . . . S8

S7 Intermediate-specific distances . . . . . . . . . . . . . . . . . . . . . . . . . . . . S9
S8 Intermediate specific locks broken between I2 and I3 . . . . . . . . . . . . . . S10
S9 Errors for plots in Figure 5 . . . . . . . . . . . . . . . . . . . . . . . . . . . . . . S11
S10 Interactions energies for STE2 and CB1 . . . . . . . . . . . . . . . . . . . . . . . S12
S11 Residues in STE2 TM7 responsible for activation . . . . . . . . . . . . . . . . S13
S12 Adaptive Sampling - 1 . . . . . . . . . . . . . . . . . . . . . . . . . . . . . . . . S14
S13 Adaptive Sampling - 2 . . . . . . . . . . . . . . . . . . . . . . . . . . . . . . . . S15
S14 Adaptive Sampling - 3 . . . . . . . . . . . . . . . . . . . . . . . . . . . . . . . . S16
S15 Implied Timescales and VAMP score plots . . . . . . . . . . . . . . . . . . . . S17
S16 Chapman Kolmogorov Test for MSM validation . . . . . . . . . . . . . . . . . . S18

### List of Tables

S1 Lipid Composition for membranes . . . . . . . . . . . . . . . . . . . . . . . . . S19
S2 Metrics for adaptive sampling. Pairs of distances between residue 1 and residue 2 for both protomers were used as distances for intra-protomer contacts. For inter-protomer contacts, distances were calculated by considering the two residues in different protomers. . . . . . . . . . . . . . . . . . . . . . . . S20
S3 Adaptive sampling roundwise data collection . . . . . . . . . . . . . . . . . . . S21
S4 Residue-level constraints for ProteinMPNN - STE2 . . . . . . . . . . . . . . . S22
S5 Residue-level constraints for ProteinMPNN - CB1 . . . . . . . . . . . . . . . . S22

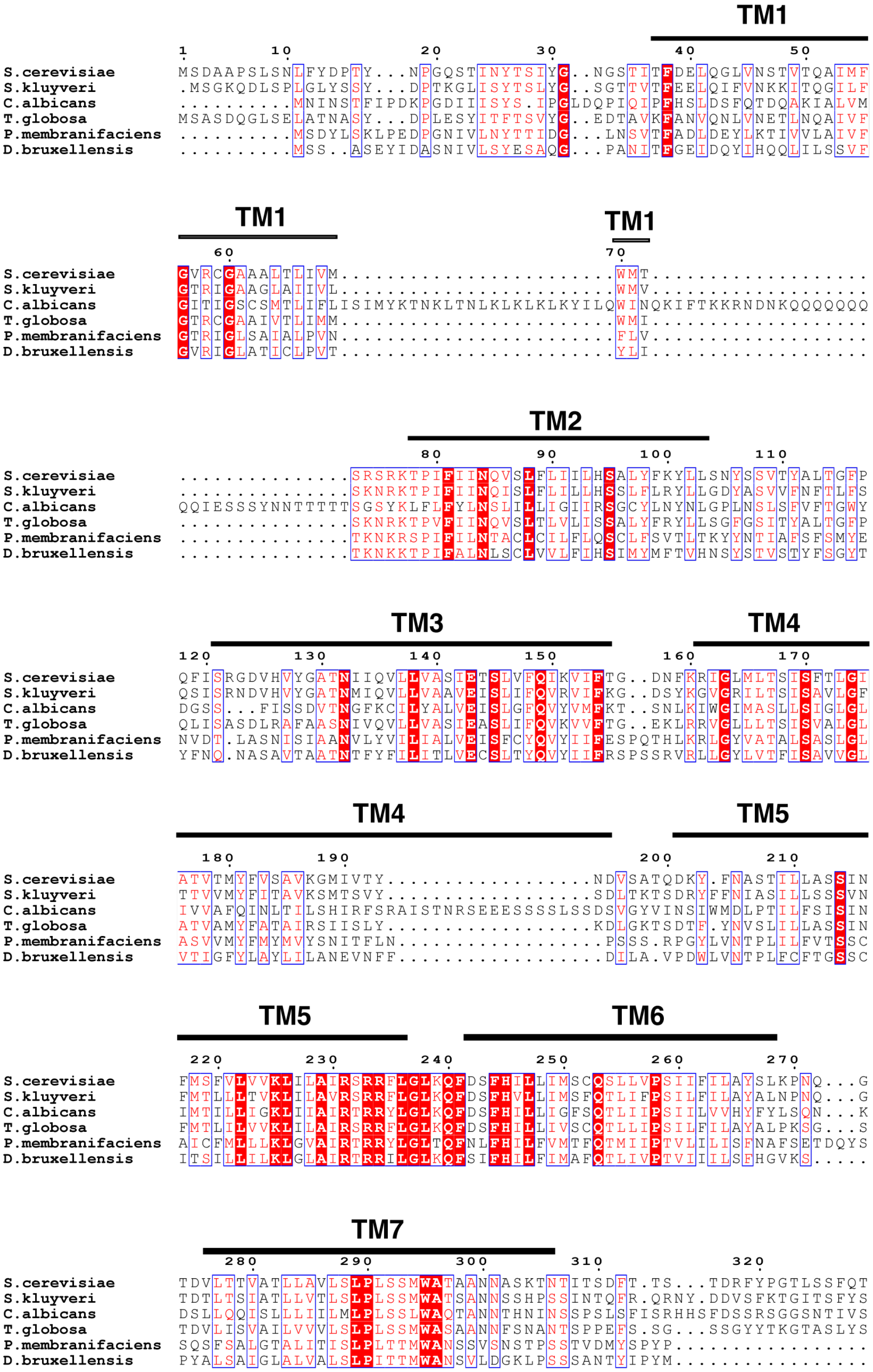

Figure S1: Multiple sequence alignments of Class D GPCRs across different fungi. Alpha-pheromone receptors were used to generate the multiple sequence alignments. Made using ESPript webserver.[1] Uniprot IDs for the sequences used: D6VTK4, P12384, A0A1D8PTB4, A0A7G3ZE52, A0A1E3NJA7, A0A871R0A3.

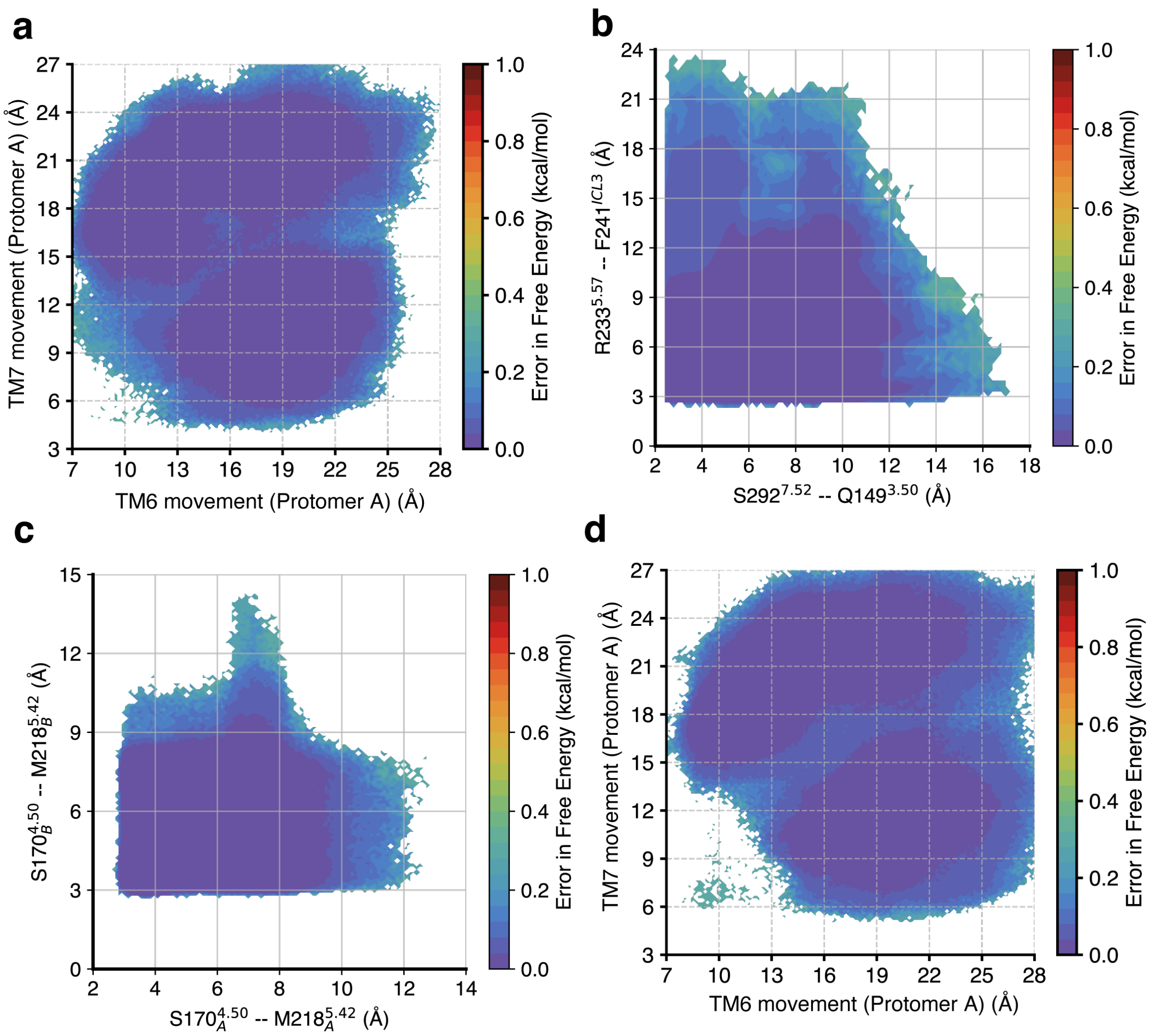

Figure S2: (a-d) Errors for free energy plots presented in Fig2, Fig3 and FigS5. (a) Error in Free Energies for (a) Fig. 2a/Fig. 5d/S5a/ (b) Fig. 2c (c) Fig. 3a (d) Fig. S5d. Errors were calculated using 200 bootstrapped MSMs, each constructed with 80% of the trajectories chosen randomly, with a lag time of 30ns.

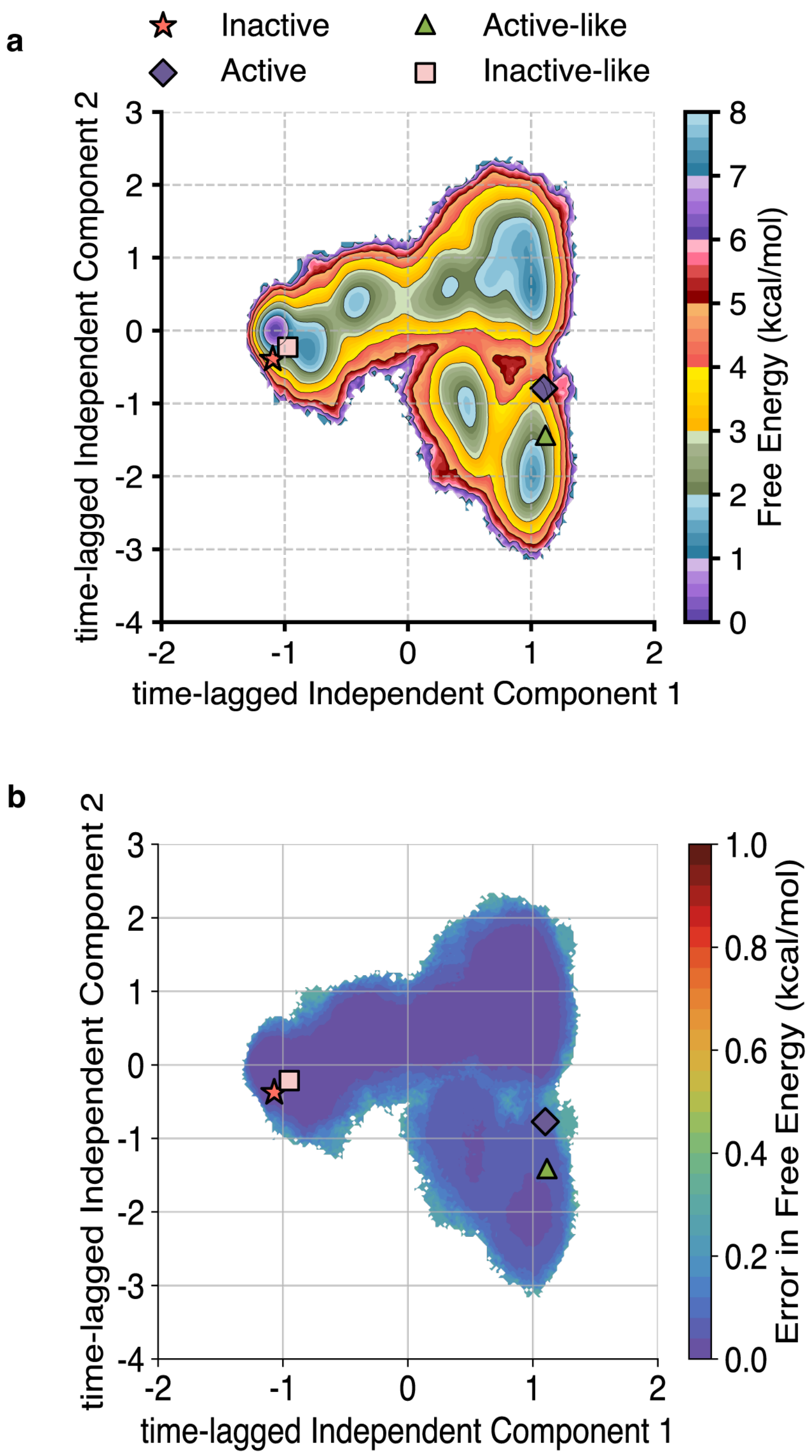

Figure S3: (a) Free Energy estimates for each simulation frame projected using time-lagged Independent Component Analysis (tICA), along the first two dimensions for the entire dataset. The slowest process (along tIC1) is activation of STE2. (b) Errors for the free energies in (a). Errors were calculated using 200 bootstrapped MSMs, each constructed with 80% of the trajectories chosen randomly, with a lag time of 30ns.

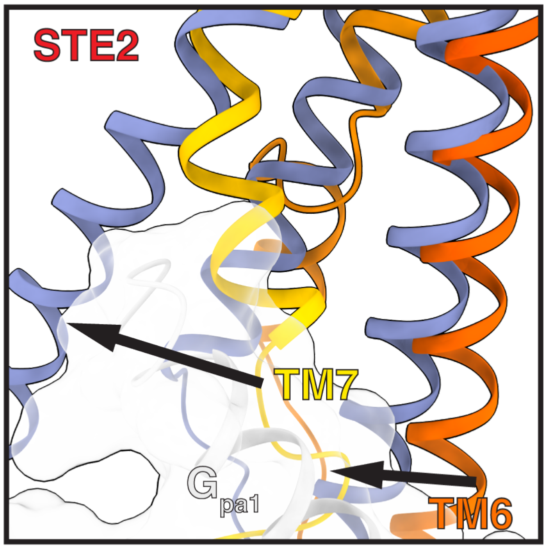

Figure S4: Inward movement of TM6 upon activation of STE2.

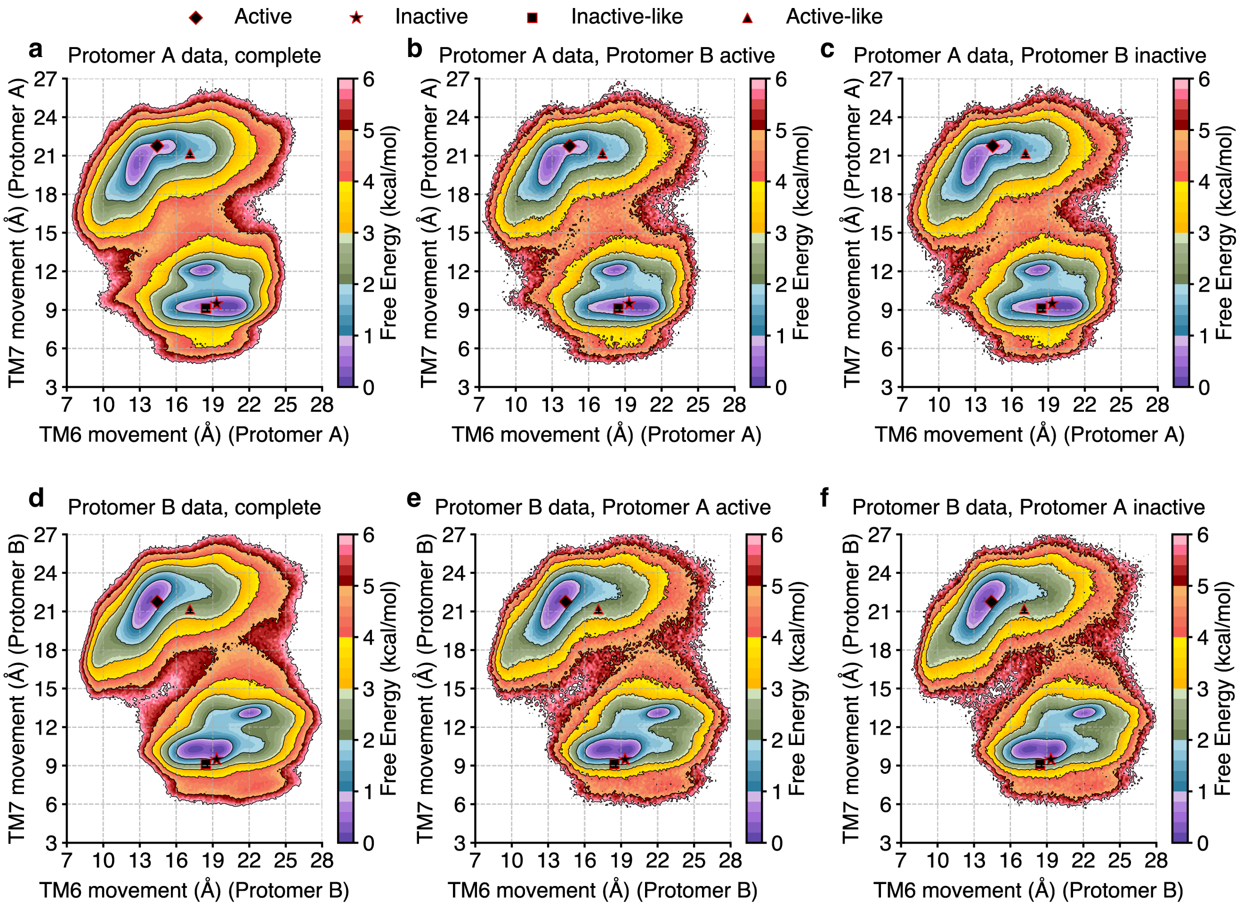

Figure S5: Free energy landscapes projection activation metrics on a monomer-specific basis. (a) TM7 movement v/s TM6 movement plotted for the complete dataset measured at protomer A. Errors for free energies plotted in Fig. S5a. (b) The same metrics as (a), but only containing data where protomer B is active. (c) The same metrics as (a), but only containing data where protomer B is inactive. (d) TM7 movement v/s TM6 movement plotted for the complete dataset measured at protomer B. Errors for free energies plotted in Fig. S5d. (e) The same metrics as (d), but only containing data where protomer A is active. (f) The same metrics as (d), but only containing data where protomer A is inactive. From the plots, it can be concluded that the one monomer being in an active/inactive state does not affect the activity of the other monomer.

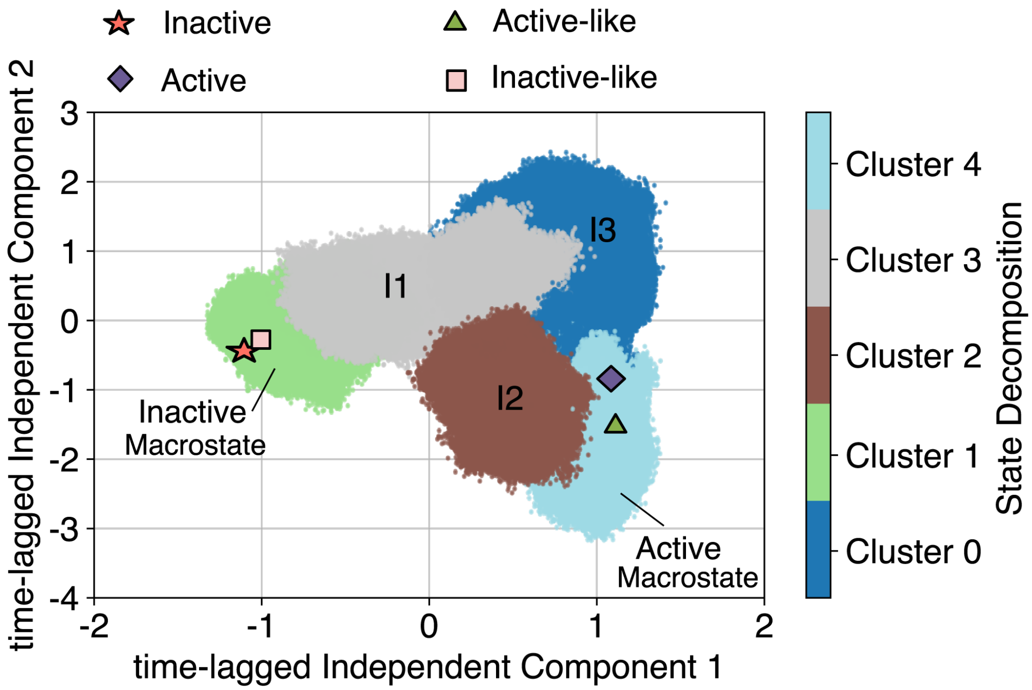

Figure S6: Macrostate decomposition projected on the slowest kinetic dimensions, tIC1 and tIC2. Inactive-like and active-like structures are identified to be in the same macrostate as inactive and active structures, respectively. Each macrostate (inactive, active, I1, I2, I3) is labelled. Comparisons with the free energies plotted on the tIC space can be made by referring to Fig. S3.

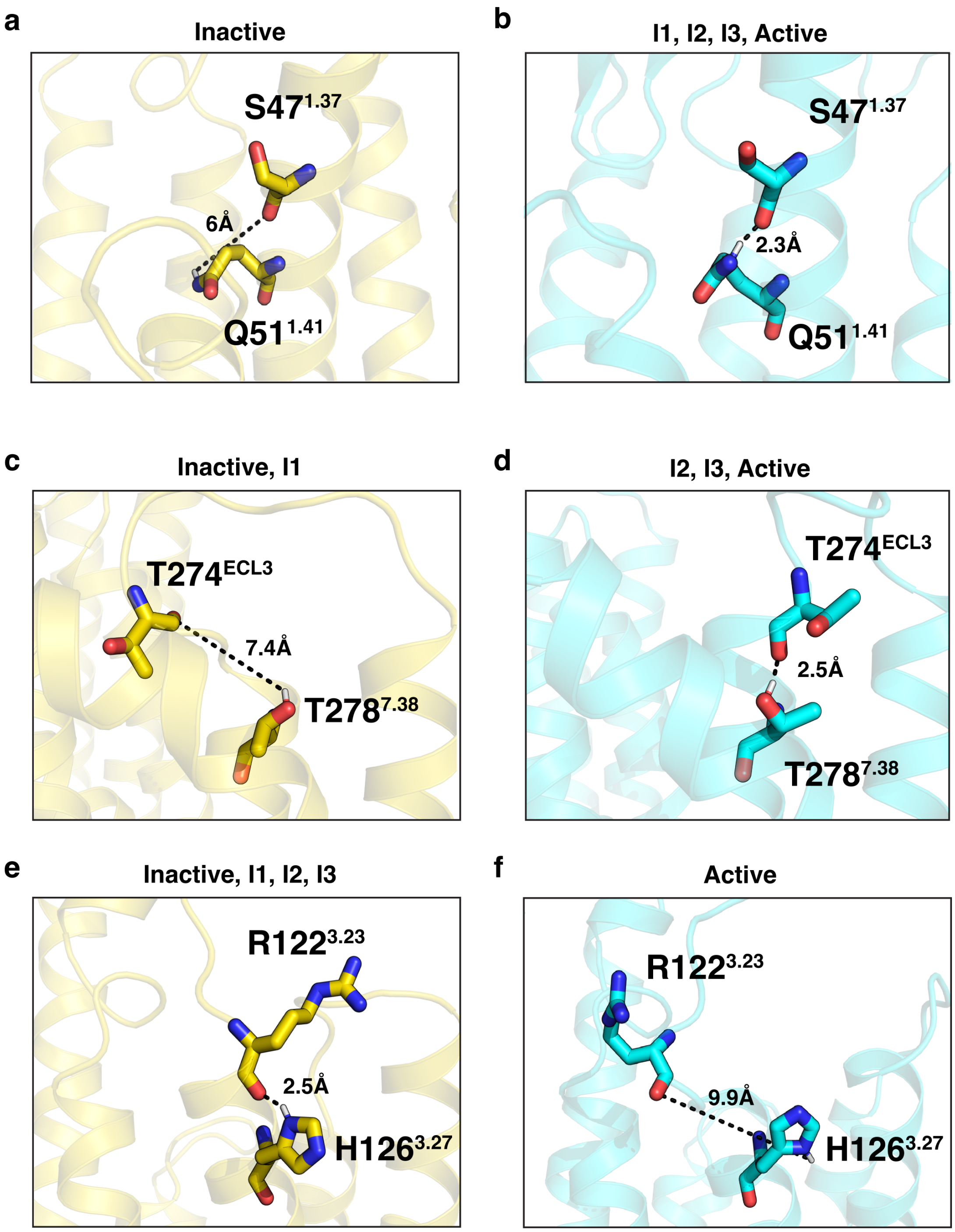

Figure S7: Distances specific to intermediate states identified in the study. (a, b) Inactive-specific molecular lock between S47$^{1.37}$ and Q51$^{1.41}$ is formed on transition to I1, and is retained along the active pathway thereafter. The probability density distribution for this distance is plotted in the main text (Fig. 4b). (c, d) Inactive and I1-specific molecular lock between T274$^{ECL3}$ and T278$^{7.38}$ is formed on transition to I2/I3, and is retained along the active pathway thereafter. The probability density distribution for this distance is plotted in the main text (Fig. 4c). (e, f) Inactive, I1, I2 and I3 specific molecular lock between R122$^{3.23}$ and H126$^{3.27}$ is broken on transition to Active. Probability density distribution for this distance is plotted in the main text (Fig. 4d).

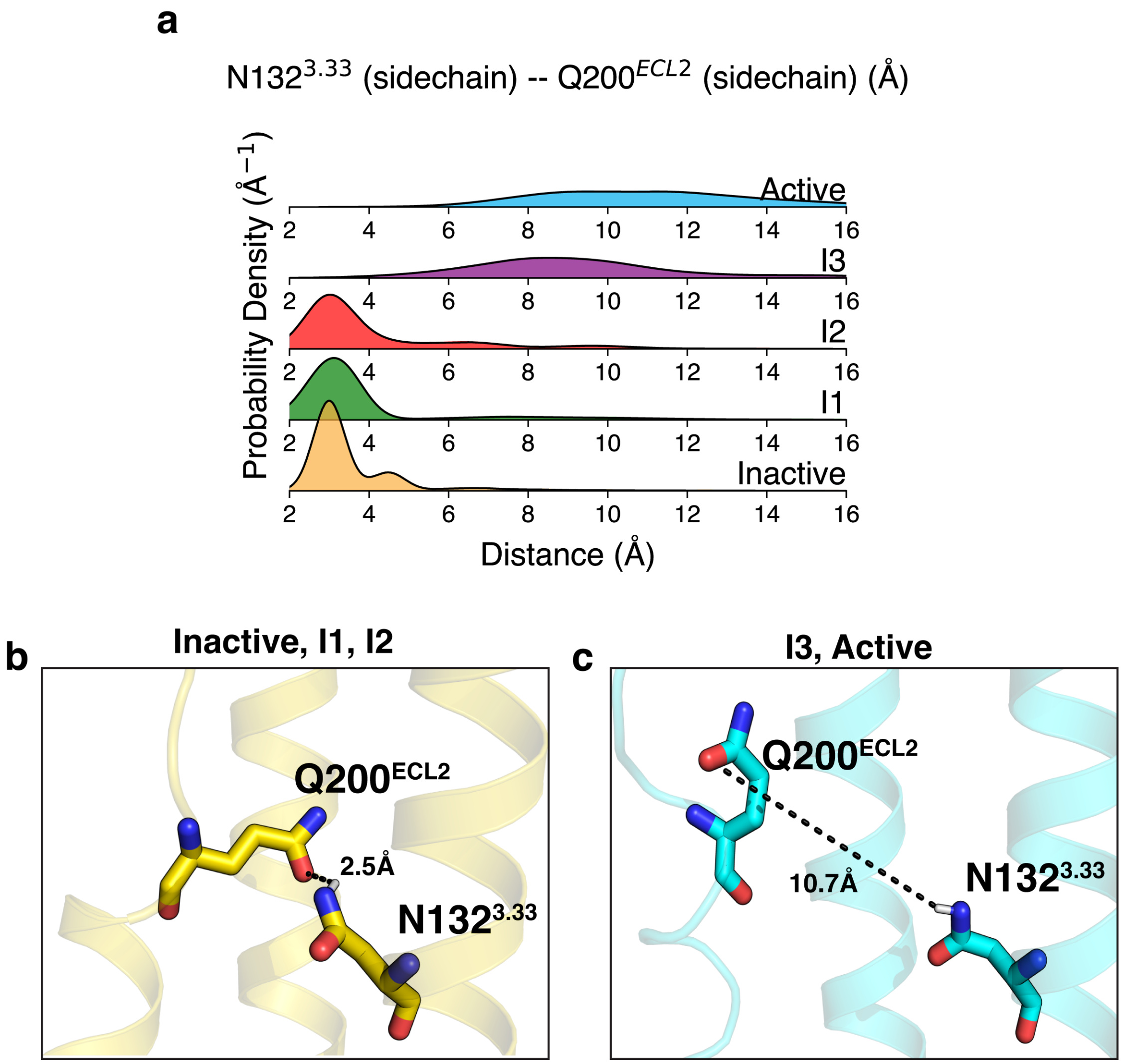

Figure S8: (a) Intermediate-state specific probability density distribution for the distance between N132$^{3.33}$ and Q200$^{ECL2}$. (b, c) Inactive, I1, I2-specific molecular lock between N132$^{3.33}$ and Q200$^{ECL2}$ is formed on transition to I3, and remains broken thereafter.

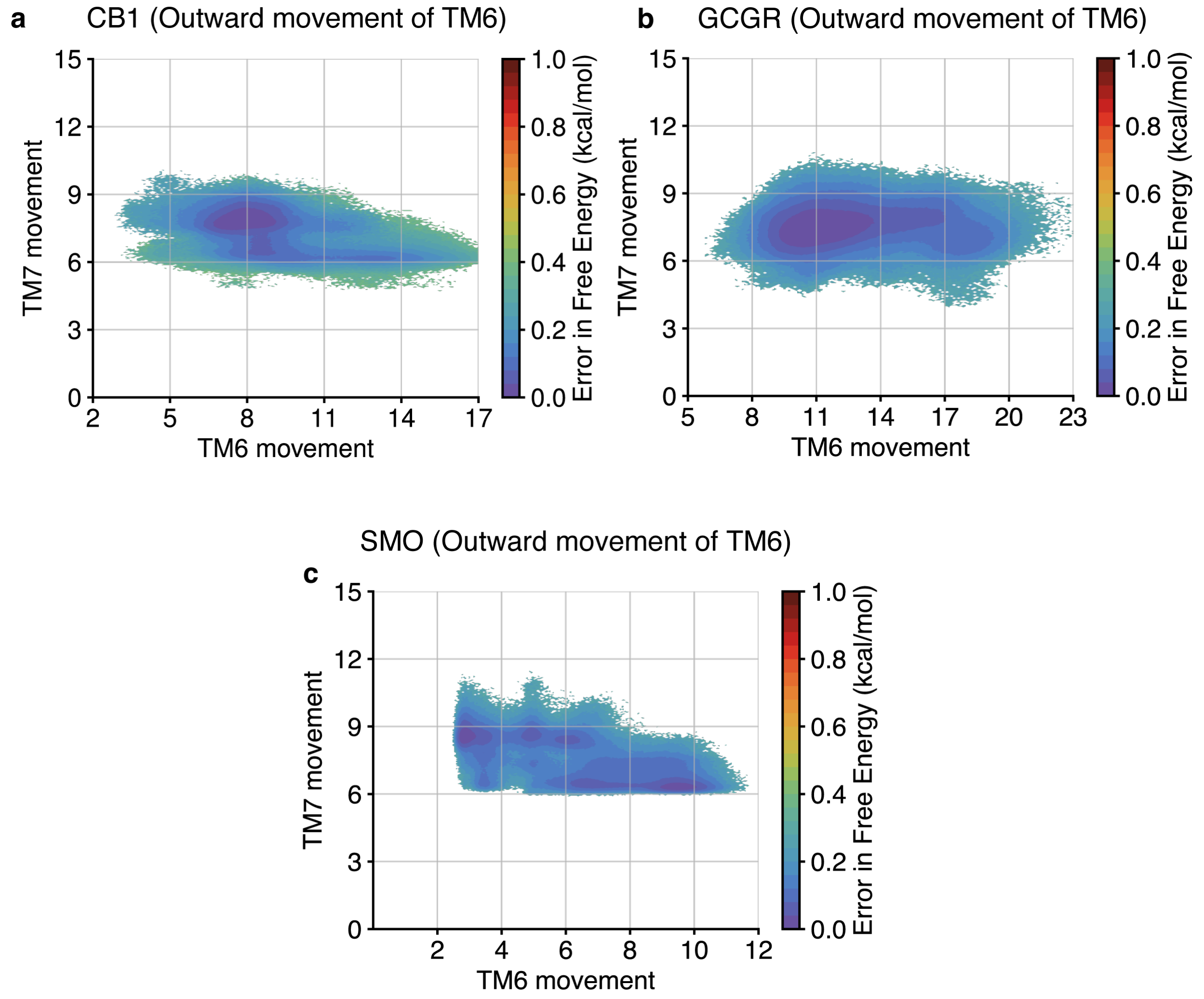

Figure S9: (a-c) Errors for free energy plots presented in Fig5. (a) Error in Free Energies for (a) Fig. 5b, (b) Fig. 5d, (c) Fig. 5f. Errors were calculated using 200 bootstrapped MSMs, each constructed with 80% of the trajectories chosen randomly, with a lagtime of 30ns.

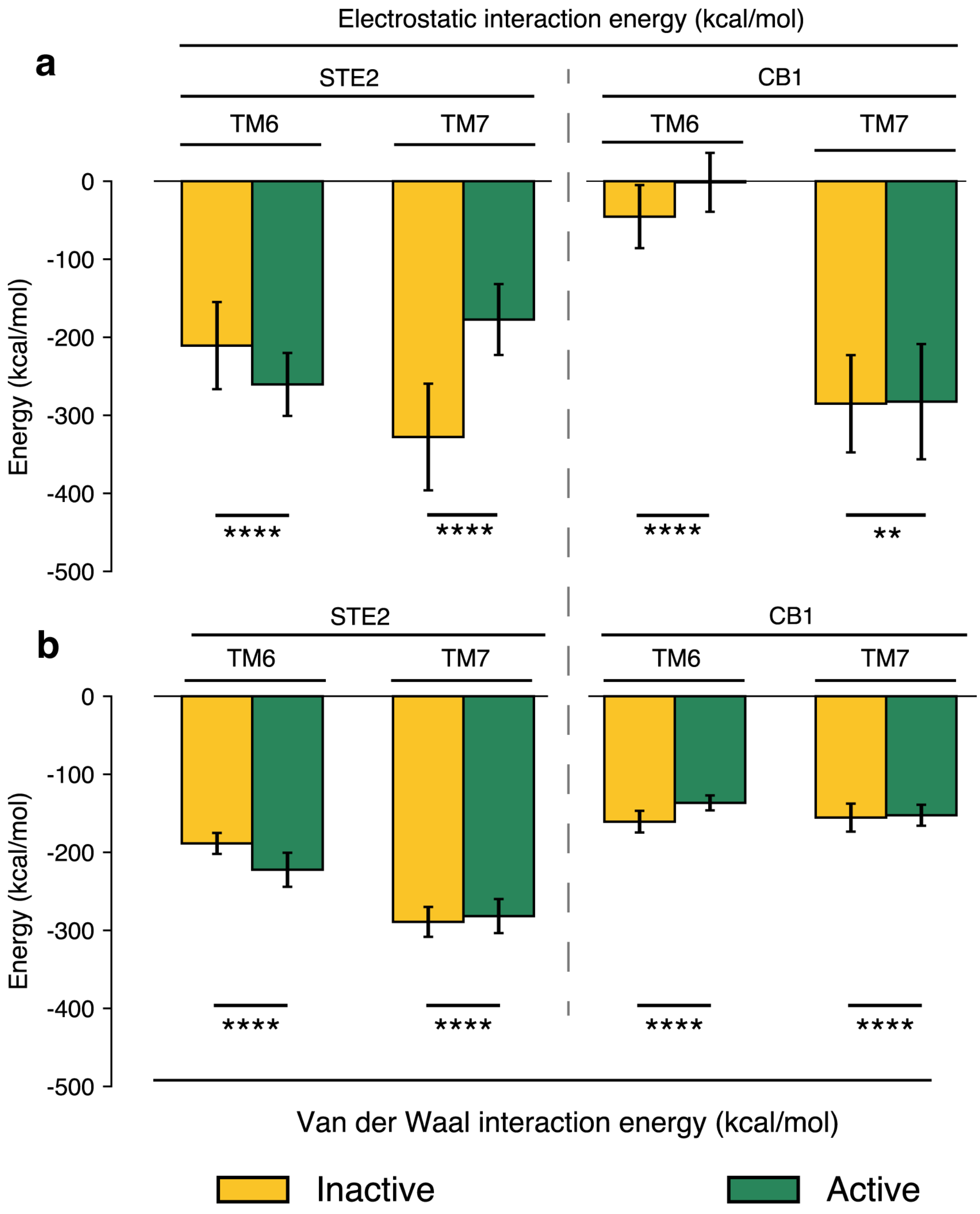

Figure S10: Interaction energies for STE2 and CB1. (a) Electrostatic energy calculated between TM6 and TM7 for STE2 (left) and CB1(right), for inactive(yellow) and active (green) conformations. (b) Same as (a), but for Van der Waals energies. p-values were calculated with a t-test with Welch's correction. P values: Inactive vs Active, TM6, STE2, electrostatic: $< 10^{-10}$; Inactive vs Active, TM7, STE2, electrostatic: $< 10^{-10}$; Inactive vs Active, TM6, CB1, electrostatic: $< 10^{-10}$; Inactive vs Active, TM7, CB1, electrostatic: 0.0086; Inactive vs Active, TM6, STE2, Van der Waal: $< 10^{-10}$; Inactive vs Active, TM7, STE2, Van der Waal: $< 10^{-10}$; Inactive vs Active, TM6, CB1, Van der Waal: $< 10^{-10}$; Inactive vs Active, TM7, CB1, Van der Waal: $< 10^{-10}$; key: Not significant (ns) $P > 0.05$, $*P \leq 0.05$, $**P \leq 0.01$, $***P \leq 0.001$, and $****P \leq 0.0001$, N=10000 for all experiments.

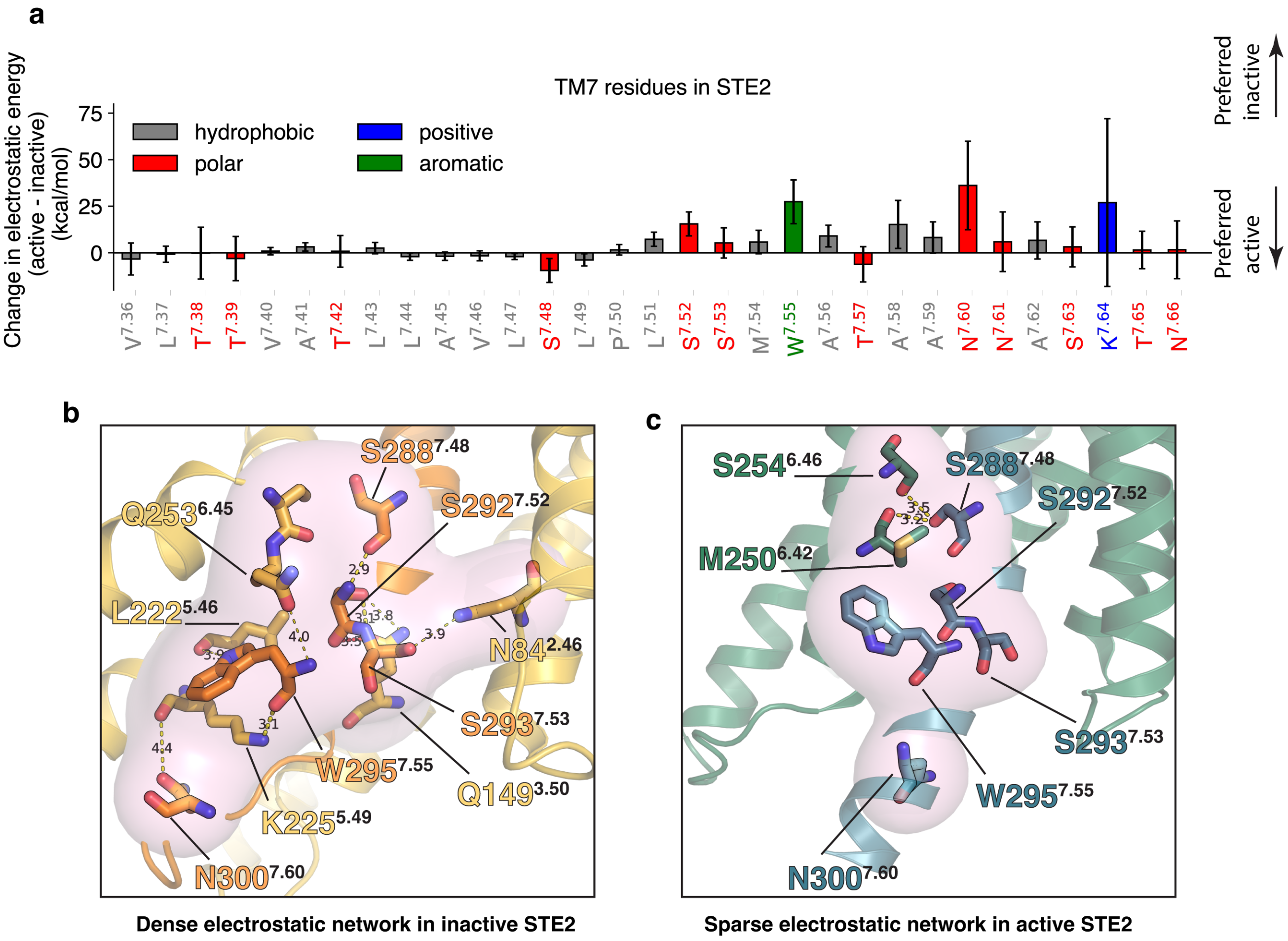

Figure S11: Residues in STE2 TM7 responsible for activation. (a) Difference in electrostatic energies (active - inactive) for each residue in TM7. Positive values imply electrostatic interactions in the inactive state. (b) A dense electrostatic network observed in TM7 of the STE2 inactive state, involving TM6, TM5, and TM3. (c) A sparse electrostatic network is observed in the STE2 active state, owing to the outward movement of TM7.

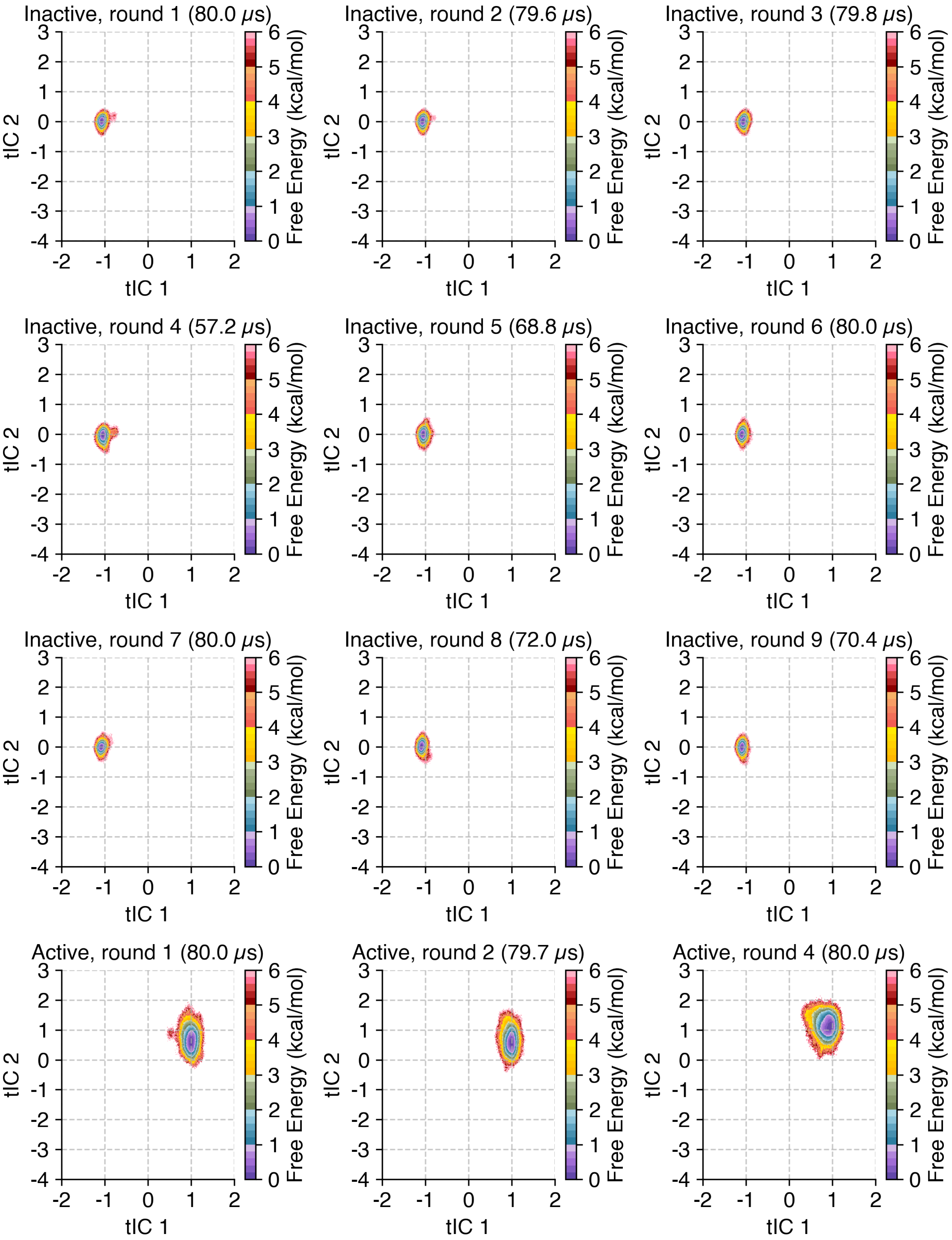

Figure S12: Adaptive Sampling performed for simulations. Plots represent the fraction of the total data collected for a specific round of simulations, grouped according to the starting point used to simulate them. Plots represent the data projected on the first two (slowest) components of the tIC space defined in Fig. S3. The title of the plot represents the starting conformation for the particular dataset, followed by the round number, and the amount of MD simulations performed for that round.

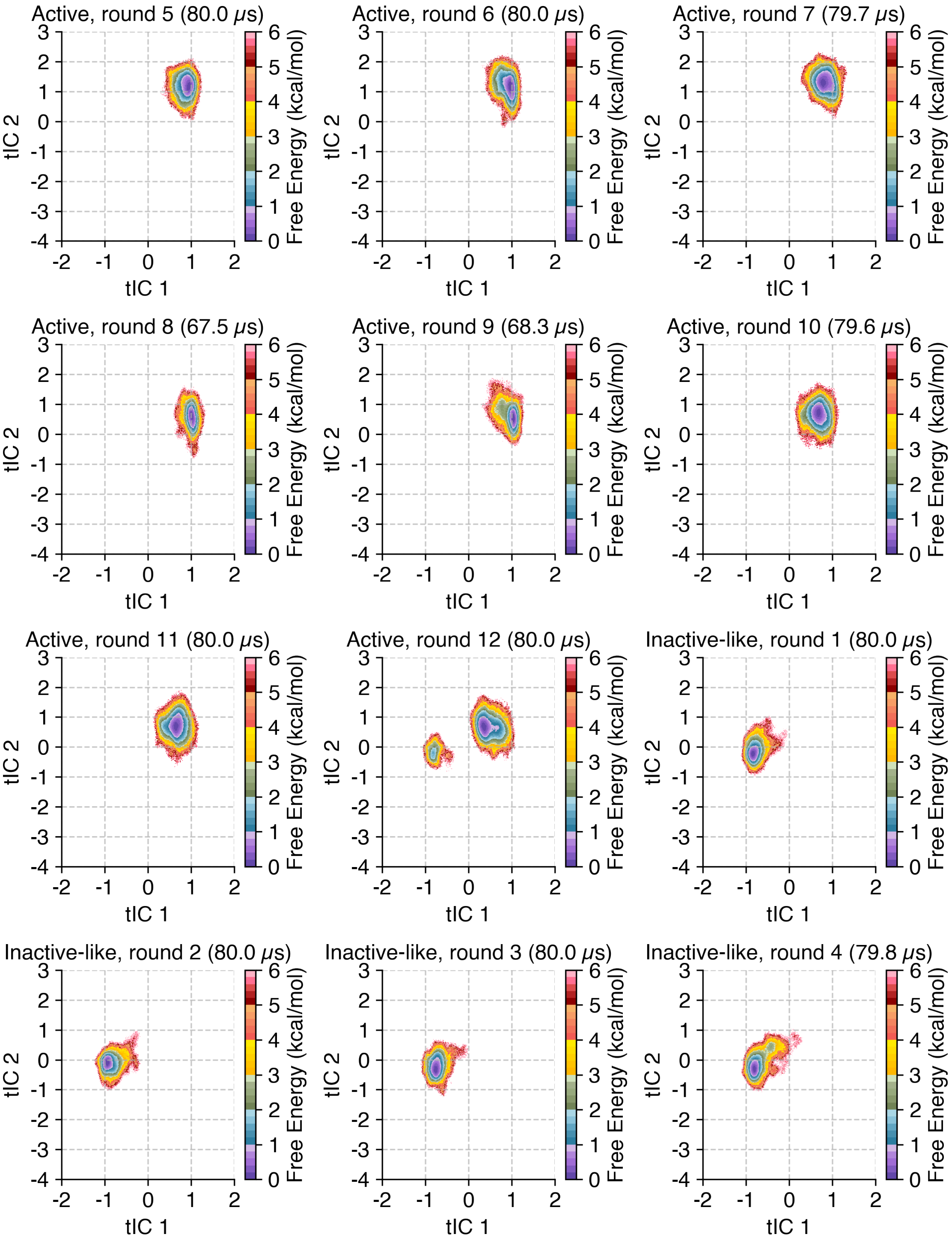

Figure S13: Adaptive Sampling performed for simulations, continued. Plots represent the fraction of the total data collected for a specific round of simulations, grouped according to the starting point used to simulate them. Plots represent the data projected on the first two (slowest) components of the tIC space defined in Fig. S3. The title of the plot represents the starting conformation for the particular dataset, followed by the round number, and the amount of MD simulations performed for that round.

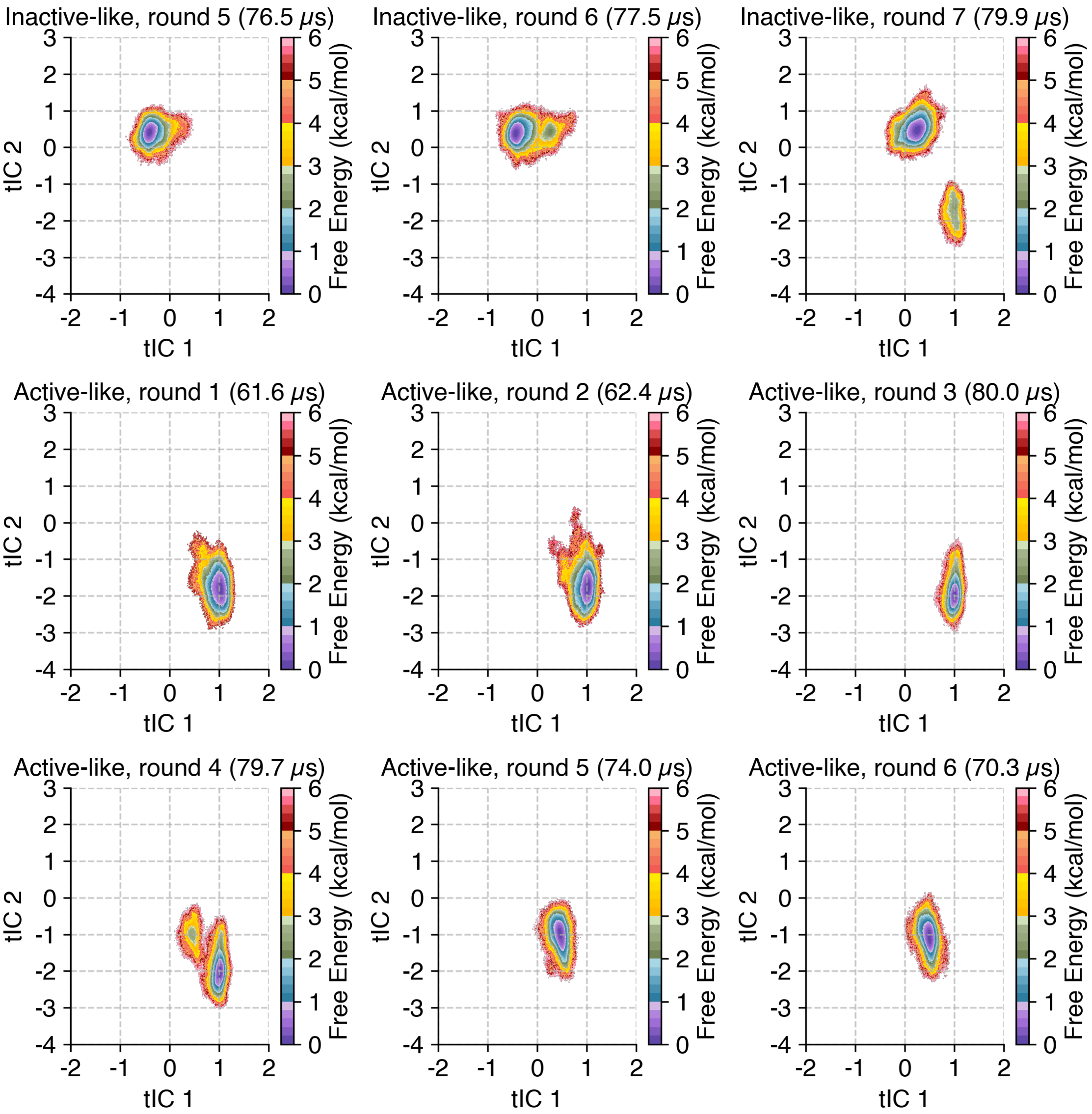

Figure S14: Adaptive Sampling performed for simulations, continued. Plots represent the fraction of the total data collected for a specific round of simulations, grouped according to the starting point used to simulate them. Plots represent the data projected on the first two (slowest) components of the tIC space defined in Fig. S3. The title of the plot represents the starting conformation for the particular dataset, followed by the round number, and the amount of MD simulations performed for that round.

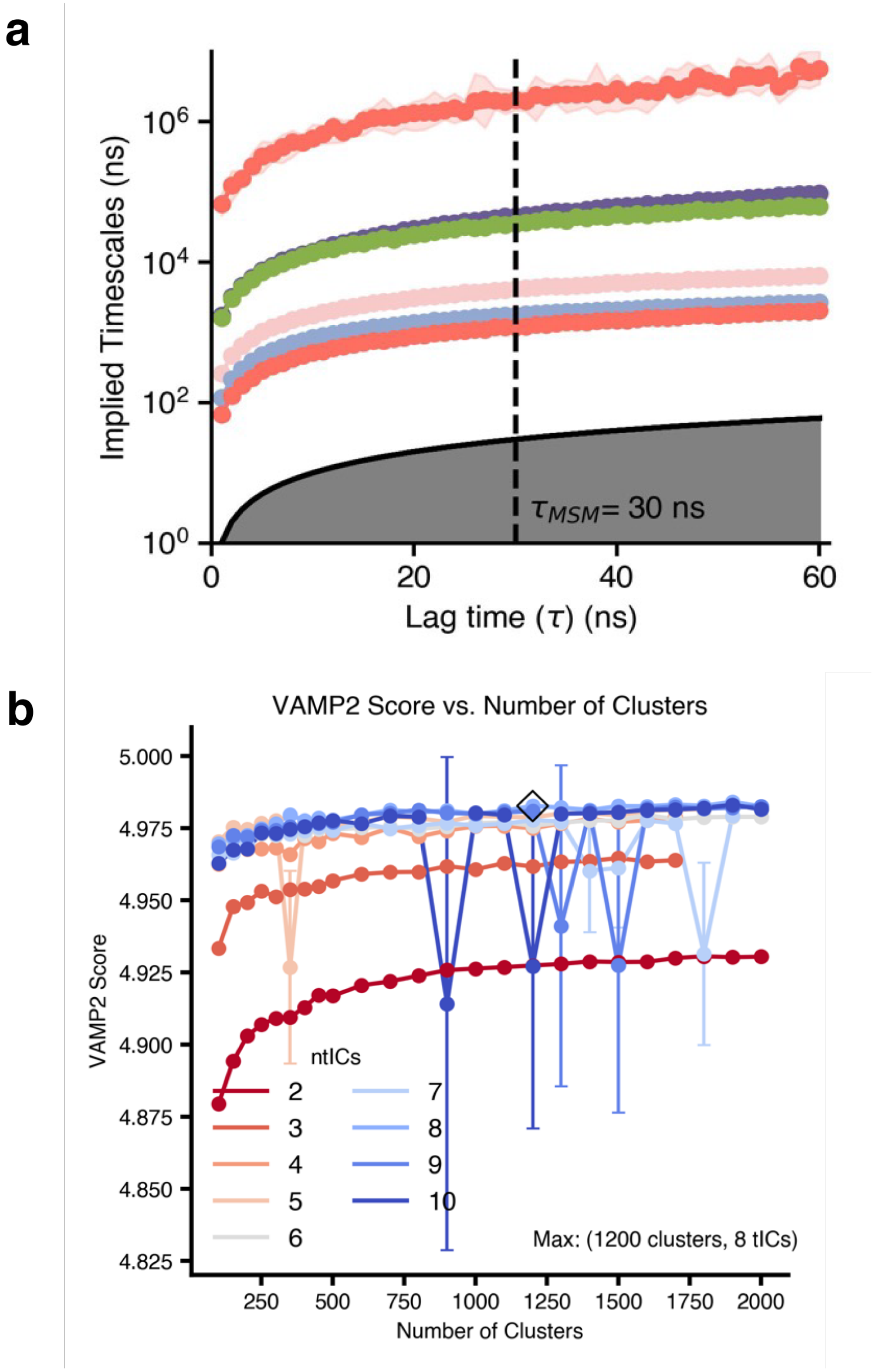

Figure S15: Implied Timescales and VAMP score plots for the constructed MSM. (a) Implied Timescales plot v/s MSM lagtime for the Markov State Model. A lag time of 30 ns was chosen for the construction of the model. (b) VAMPscore as a function of the number of states and number of tIC components. 1200 clusters and 8 tICs were chosen for the input to the state decomposition (shown as a black diamond).

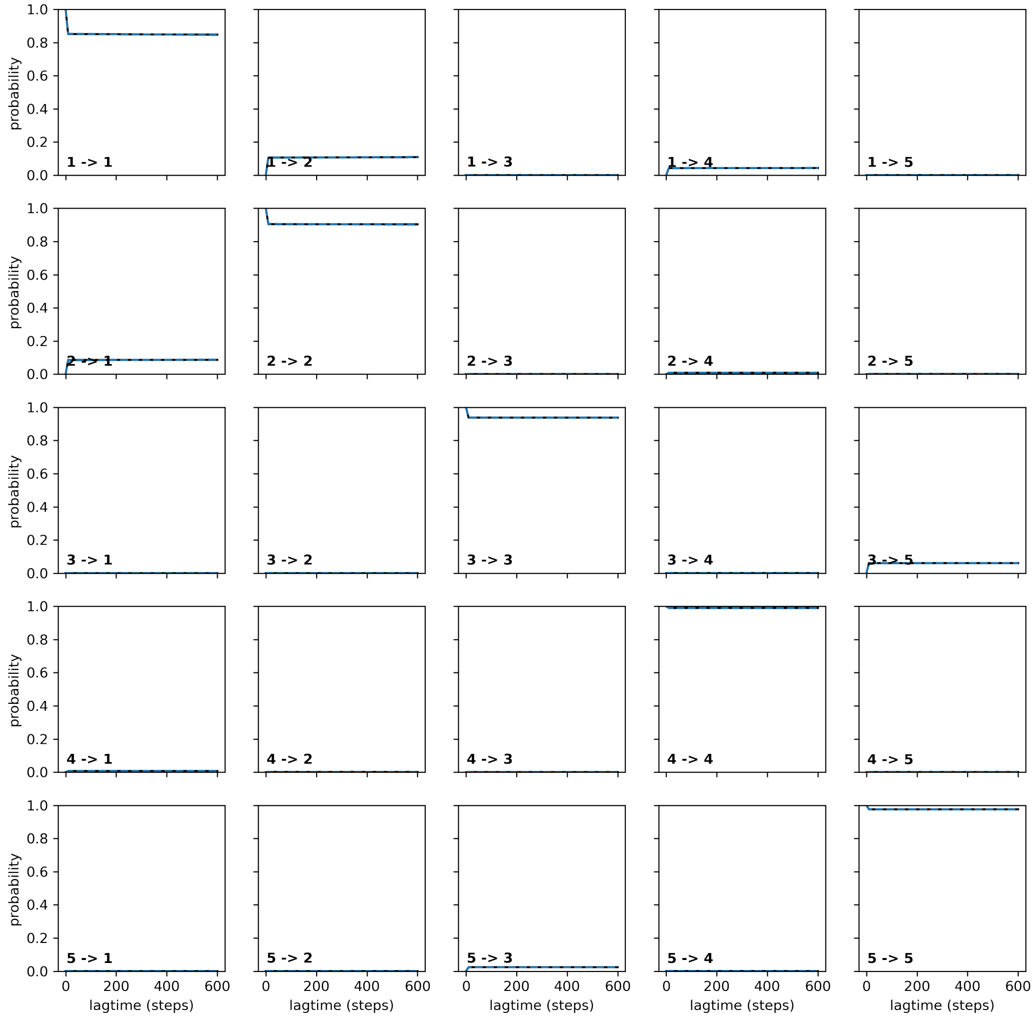

Figure S16: Chapman Kolmogorov Test for MSM validation. 5 macrostates were used.

Table S1: Lipid Composition for membranes

| **Lipid (Full Name)** | % |
|---|---|
| Ergosterol | 4 |
| Ceramide 241 + 2-$\beta$-D-mannose + 2-$\beta$-D-glucose | 2 |
| Ceramide 241 + 2-$\beta$-D-mannose + 1-$\beta$-D-glucose (2-neuphos) + 1-$\alpha$-D-glucose | 4 |
| Ceramide 241 + 2-$\beta$-D-mannose | 12 |
| 1-Palmitoyl-2-hydroxy-sn-glycero-3-phosphocholine (PYPC) | 4 |
| 1-Oleoyl-2-palmitoyl-sn-glycero-3-phosphocholine (YOPC) | 17 |
| 1-Palmitoyl-2-oleoyl-sn-glycero-3-phosphocholine (POPC) | 4 |
| 1,2-Dioleoyl-sn-glycero-3-phosphocholine (DOPC) | 2 |
| 1,2-Dioleoyl-sn-glycero-3-phosphoethanolamine (DYPE) | 3 |
| 1-Palmitoyl-2-hydroxy-sn-glycero-3-phosphoethanolamine (PYPE) | 1 |
| 1-Oleoyl-2-palmitoyl-sn-glycero-3-phosphoethanolamine (YOPE) | 7 |
| 1-Palmitoyl-2-oleoyl-sn-glycero-3-phosphoethanolamine (POPE) | 3 |
| 1-Palmitoyl-2-oleoyl-sn-glycero-3-phospho-L-serine (POPS) | 2 |
| 1-Palmitoyl-2-linoleoyl-sn-glycero-3-phospho-L-serine (PLPS) | 3 |
| 1-Palmitoyl-2-oleoyl-sn-glycero-3-phosphoinositol (POPI) | 14 |
| 1-Palmitoyl-2-hydroxy-sn-glycero-3-phosphoinositol (PYPI) | 5 |
| 1-Palmitoyl-2-linoleoyl-sn-glycero-3-phosphoinositol (PLPI) | 3 |
| 1,2-Dioleoyl-sn-glycero-3-phosphocholine (DYPC) | 10 |
| **Total** | **100** |

Table S2: Metrics for adaptive sampling. Pairs of distances between residue 1 and residue 2 for both protomers were used as distances for intra-protomer contacts. For inter-protomer contacts, distances were calculated by considering the two residues in different protomers.

| Residue 1 | Residue 2 | Residue 1 | Residue 2 | Residue 1 | Residue 2 | Contact type |
|---|---|---|---|---|---|---|
| F12 | G31 | S87 | S141 | N216 | L255 | intra-protomer |
| N18 | Y26 | Y98 | G188 | S219 | S254 | intra-protomer |
| P19 | I24 | Y106 | I120 | F220 | L255 | intra-protomer |
| G20 | D39 | Y106 | V196 | K225 | I230 | intra-protomer |
| Q21 | S107 | Q118 | A185 | K225 | R233 | intra-protomer |
| Y26 | S34 | V125 | Y193 | K225 | R234 | intra-protomer |
| T27 | N32 | H126 | Y193 | S232 | N301 | intra-protomer |
| N46 | V257 | S141 | L284 | R233 | L238 | intra-protomer |
| N46 | K269 | S145 | S288 | R233 | F241 | intra-protomer |
| S47 | L90 | Q149 | S292 | L238 | L287 | intra-protomer |
| S47 | G273 | I153 | A296 | Q240 | H245 | intra-protomer |
| T50 | N271 | F154 | K225 | Q240 | L287 | intra-protomer |
| M54 | A265 | I162 | L210 | H245 | L283 | intra-protomer |
| M54 | L277 | I169 | K202 | I246 | W295 | intra-protomer |
| F55 | Y98 | S170 | M218 | L248 | Q253 | intra-protomer |
| A62 | K77 | T179 | D195 | L248 | W295 | intra-protomer |
| W70 | R76 | K187 | Y203 | M250 | G273 | intra-protomer |
| W70 | Q85 | F204 | S254 | M250 | S288 | intra-protomer |
| T72 | F235 | L211 | I246 | Q253 | L291 | intra-protomer |
| I80 | S243 | A212 | L247 | V257 | Y266 | intra-protomer |
| P258 | A281 | L284 | S141 | S292 | Q149 | intra-protomer |
| A265 | T274 | S288 | S145 | A296 | I153 | intra-protomer |
| Y17 | T110 | L64 | G56 | S108 | S34 | inter-protomer |
| P19 | Y111 | M71 | L287 | Q118 | T603 | inter-protomer |
| W295 | A63 | N301 | S293 | | | inter-protomer |

Table S3: Adaptive sampling rounds and collected simulation times. The first column indicates the starting conformational state (PDB structure), the second the adaptive sampling round, and the third the amount of data collected in that round.

| Starting Point | Round | Total data collected ($\mu$s) |
|---|---|---|
| Inactive (7QA8) | 1 | 80.0 |
| Inactive (7QA8) | 2 | 79.6 |
| Inactive (7QA8) | 3 | 79.8 |
| Inactive (7QA8) | 4 | 57.2 |
| Inactive (7QA8) | 5 | 68.8 |
| Inactive (7QA8) | 6 | 80.0 |
| Inactive (7QA8) | 7 | 80.0 |
| Inactive (7QA8) | 8 | 72.0 |
| Inactive (7QA8) | 9 | 70.4 |
| Active (7AD3) | 1 | 80.0 |
| Active (7AD3) | 2 | 79.7 |
| Active (7AD3) | 4 | 80.0 |
| Active (7AD3) | 5 | 80.0 |
| Active (7AD3) | 6 | 80.0 |
| Active (7AD3) | 7 | 79.7 |
| Active (7AD3) | 8 | 67.5 |
| Active (7AD3) | 9 | 68.3 |
| Active (7AD3) | 10 | 79.6 |
| Active (7AD3) | 11 | 80.0 |
| Active (7AD3) | 12 | 80.0 |
| Inactive-like (7QBC) | 1 | 80.0 |
| Inactive-like (7QBC) | 2 | 80.0 |
| Inactive-like (7QBC) | 3 | 80.0 |
| Inactive-like (7QBC) | 4 | 79.8 |
| Inactive-like (7QBC) | 5 | 76.5 |
| Inactive-like (7QBC) | 6 | 77.5 |
| Inactive-like (7QBC) | 7 | 79.9 |
| Active-like (7QBI) | 1 | 61.6 |
| Active-like (7QBI) | 2 | 62.4 |
| Active-like (7QBI) | 3 | 80.0 |
| Active-like (7QBI) | 4 | 79.7 |
| Active-like (7QBI) | 5 | 74.0 |
| Active-like (7QBI) | 6 | 70.3 |

Table S4: Residue-level constraints applied during ProteinMPNN sequence generation in STE2. The same positions were restrained in both monomers.

| | | | | |
|---|---|---|---|---|
| G31 | F38 | G56 | G60 | F81 |
| N84 | L88 | S95 | N132 | L138 |
| E143 | S145 | Q149 | F154 | G163 |
| S170 | G174 | S214 | L222 | K225 |
| L226 | A229 | R231 | R233 | R234 |
| L236 | G237 | L238 | Q240 | F241 |
| F244 | H245 | L247 | Q253 | P258 |
| L289 | P290 | W295 | A296 | |

Table S5: Residue-level constraints applied during ProteinMPNN sequence generation for CB1.

| | | | | |
|---|---|---|---|---|
| C139 | D163 | V168 | L209 | I218 |
| H219 | R220 | A236 | V246 | N256 |
| V291 | H302 | S303 | K343 | L345 |
| L352 | C355 | W356 | P358 | N389 |
| N393 | P394 | Y397 | | |
